# *Eomes* restricts *Brachyury* functions at the onset of mammalian gastrulation

**DOI:** 10.1101/2023.01.27.525830

**Authors:** Katrin M. Schüle, Jelena Weckerle, Simone Probst, Alexandra E. Wehmeyer, Lea Zissel, Chiara M. Schröder, Mehmet Tekman, Gwang-Jin Kim, Inga-Marie Schlägl, Sagar, Sebastian J. Arnold

## Abstract

Mammalian specification of mesoderm and definitive endoderm (DE) is instructed by the two related Tbx transcription factors (TFs) *Eomesodermin* (*Eomes*) and *Brachyury* sharing partially redundant functions. Gross differences of mutant embryonic phenotypes suggest specific functions of each TF. To date, the molecular details of separated lineage-specific gene-regulation by *Eomes* and *Brachyury* remain poorly understood. Here, we combine embryonic and stem cell-based analyses to delineate the non-overlapping, lineage-specific transcriptional activities. On a genome-wide scale binding of both TFs overlaps at promoters of target genes, but shows specificity for distal enhancer regions, that is conferred by differences in Tbx DNA-binding motifs. The unique binding to enhancer sites instructs the specification of anterior mesoderm (AM) and DE by *Eomes* and caudal mesoderm by *Brachyury*. Remarkably, EOMES antagonizes BRACHYURY gene-regulatory functions in co-expressing cells during early gastrulation to ensure the proper sequence of early AM and DE lineage specification followed by posterior mesoderm derivatives.

**Highlights:**

- Detailed comparative analysis of the two critical developmental regulators *Eomes* and *Brachyury* in mouse embryos and differentiating embryonic stem cells
- Tbx factors EOMES and BRACHYURY control distinct gene programs to specify different mesoderm and endoderm subsets
- Program specificity is conferred by binding to non-overlapping enhancers with distinct binding motifs
- EOMES restricts the activities of BRACHYURY thus ensuring the proper sequence of mesoderm and endoderm lineage specification

## Introduction

During mouse gastrulation pluripotent epiblast cells become lineage specified to form the three primary germ layers (neuro-) ectoderm (NE), mesoderm and definitive endoderm (DE). Early anterior mesoderm (AM) and DE specification are intrinsically linked to morphogenesis of the emerging mesoderm layer when cells from the posterior epiblast ingress through the primitive streak (PS) thus establishing a third cell layer containing both mesoderm and DE (mesoderm and endoderm, ME) progenitors. The cells that remain in the anterior epiblast give rise to NE^1^. Among the earliest genes that mark the formation of the PS and onset of gastrulation around embryonic day 6.5 (E6.5) are the Tbx transcription factors *Eomesodermin* (*Eomes*) and *Brachyury* (*T, hereafter also referred to as Bra*) ^2-4^. In the early gastrulating embryo *Eomes* is expressed in all cells on the posterior half of the epiblast that mostly contribute to anterior cell types leaving the PS before E7.5, including cardiac mesoderm and DE lineages ^5-8^. According to its expression during gastrulation initiation the functional deletion of *Eomes* from the epiblast results in early and severe gastrulation defects and the complete absence of AM and DE cell types ^5,6,9^. In contrast, during the first day of gastrulation (E6.5-E7.5) *Brachyury* expression is refined to the PS and fulfills functions during specification of extraembryonic and axial midline mesoderm ^3,10,11^. Following a first phase of gastrulation, in which mesoderm is formed by migration of epiblast cells through the elongated PS, *Brachyury* remains expressed in the neuromesodermal progenitors (NMPs) and the tailbud that support the posterior extension of the embryonic body axis during the second phase of gastrulation ^12-14^. Accordingly, *Brachyury* (*T*^2J/2J^) mutant embryos exhibit disrupted mesoderm and somite formation caudal to the seven most cranial somites, and an overall paucity of mesoderm ^2,12,15^. A failure to properly form the allantois and thus impaired chorio-allantoic fusion in *T*^2J/2J^ embryos leads to lethality between E9.5 and E10.5 ^12,16,17^.

Despite the obvious phenotypic differences of *Eomes* and *Brachyury* mutants, previous studies suggested partially redundant molecular functions that are reflected in overlapping genomic binding motifs, chromatin binding patterns, and the regulation of similar sets of target genes ^18-20^. This functional overlap is further exemplified by studies showing that the specification of all mesoderm or DE lineages relies on the activities of either of these two Tbx factors ^18,19,21^. Accordingly, the simultaneous deletion of *Eomes* and *Brachyury* activities (referred to as double KO (dKO) throughout this paper) completely abrogates the potential to form ME derivatives from pluripotent cells and embryos ^21^. Instead, dKO cells retain prolonged pluripotency and eventually differentiate to anterior epiblast/NE derivatives by the activation of genetic programs that are normally repressed by ME promoting TFs including *Eomes* and *Brachyury* and downstream transcription factors ^21^. Also this study observed EOMES and BRACHYURY binding at common genomic binding sites to initiate transcriptional activation of overlapping sets of ME target genes, similar to previous reports ^18-21^. Thus, the specific transcriptional targets that guide different genetic programs, and the molecular basis that confers specificity of these Tbx factors remain unresolved. Since previous studies did not reveal distinguishing T-box binding consensus motifs for different Tbx factors at endogenous target gene sites ^18,22,23^, additional mechanisms were suggested. These included differences in spatio-temporal expression patterns of the individual Tbx factors, which could impact on the chromatin permissiveness of target genes and thus influence transcriptional responses during embryonic development ^24-26^. Other concepts proposed the cooperativity of Tbx with other transcription factors ^23^, such as co-binding of *XBra* with *XSmad1* during mesoderm formation in *Xenopus* ^27^. Similarly, another study suggested additional signaling cues including NODAL or BMP4 to impact on BRACHYURY chromatin binding and function in human embryonic stem cells (hESCs) during mesoderm or DE differentiation ^20^. However, a comprehensive model explaining overlapping functions and target gene specificity of early Tbx factors is yet missing. In this study, we delineate the distinct and redundant functions of *Eomes* and *Brachyury* during embryonic ME specification and provide a molecular basis underlying cell-lineage specific program regulation. *Eomes* and *Brachyury* are first co-expressed during gastrulation in mouse embryos from E6.5-E7.5. However, lineage-tracing and transcriptional profiling of mutant embryos highlight grossly non-overlapping functions. Spatio-temporal differences in expression are not accountable for distinct functions since *Brachyury* expressed from the *Eomes* gene locus (*Eomes*^Bra^) does not rescue the *Eomes*-deficient phenotype. To study the molecular details of target gene regulation by *Eomes* and *Brachyury*, we employed an embryoid body-based differentiation model of mESCs that allows for fully controlled and comparable expression of both TFs. The integration of transcriptional profiles and chromatin-binding analyses allowed to define the unique functions and binding of both TFs during differential lineage specification. While chromatin binding of EOMES and BRACHYURY is frequently overlapping in promoter regions of target genes, specific transcriptional regulation is conferred by unique binding to intronic/intergenic enhancer sites distant from the promoters. This binding specificity is achieved by non-overlapping, EOMES-and BRACHYURY-specific consensus motifs. The analysis of accessible chromatin at specifically EOMES-and BRACHYURY-bound sites indicate antagonistic effects of *Eomes* on *Brachyury* functions. In line, expression of *Brachyury* targets is repressed upon *Eomes* overexpression in embryoid bodies. We propose that this antagonism explains the embryonic observations of early BRACHYURY expression in the early PS without any obvious functions in the presence of EOMES. Collectively, this study reveals the molecular basis of divergent functions of EOMES and BRACHYURY for the activation of distinct, competing gene programs of different ME lineages.

## Results

### *Eomes* and *Brachyury* specify distinct lineages at different gastrulation stages

*Eomes* and *Brachyury* are both expressed during gastrulation in partially overlapping patterns. To investigate their expression at cellular resolution, we analyzed recently published scRNA-seq datasets of different gastrulation stage embryos ^28^ (Fig S1A-C showing all cells of embryonic days (E) E6.5-E8.25 embryos). We selected all *Eomes* and/or *Brachyury* positive cells from E7.0, E7.5, and E8.0 (including *Eomes* single positive cells, *Brachyury* single positive cells and *Eomes/Brachyury* double positive cells, expression cut-off 0.3 normalized transcript counts) and clustered cells using VarID^29^ (Figures 1A, S1A-D and S1G). Plotting the expression of *Eomes* and *Brachyury* onto the UMAP maps shows that at E7.0 *Brachyury* positive cells are contained within the *Eomes* expressing population (Figures S1D-F), such as in the PS, nascent mesoderm and anterior PS. *Eomes* single positive cells additionally show a posterior epiblast signature (Figures S1D-F), and are also found within the clusters of nascent and extraembryonic mesoderm (ExM) (Figures S1D-F and 1A-C). At E7.5, first *Brachyury* single positive cells are detected that reside in the newly formed clusters of caudal epiblast and posterior mesoderm (PM) as well as in more differentiated tissues, such as the node and notochord, that already have downregulated *Eomes* expression (Figure 1A-C). At E7.5, *Eomes*-single positive cells are found in the posterior epiblast and in the DE cluster (Figure 1B). By E8.0 *Brachyury* expression is widespread while *Eomes* expression is grossly downregulated^4^ and almost no *Eomes* positive cells are found on the UMAP maps, except for the visceral endoderm (VE) (Figures S1G-I). These expression dynamics indicate a switch from an early *Eomes*-associated phase of gastrulation from E6.5-E7.5 to a *Brachyury*-associated state of gastrulation following E7.5.

**Figure 1.**
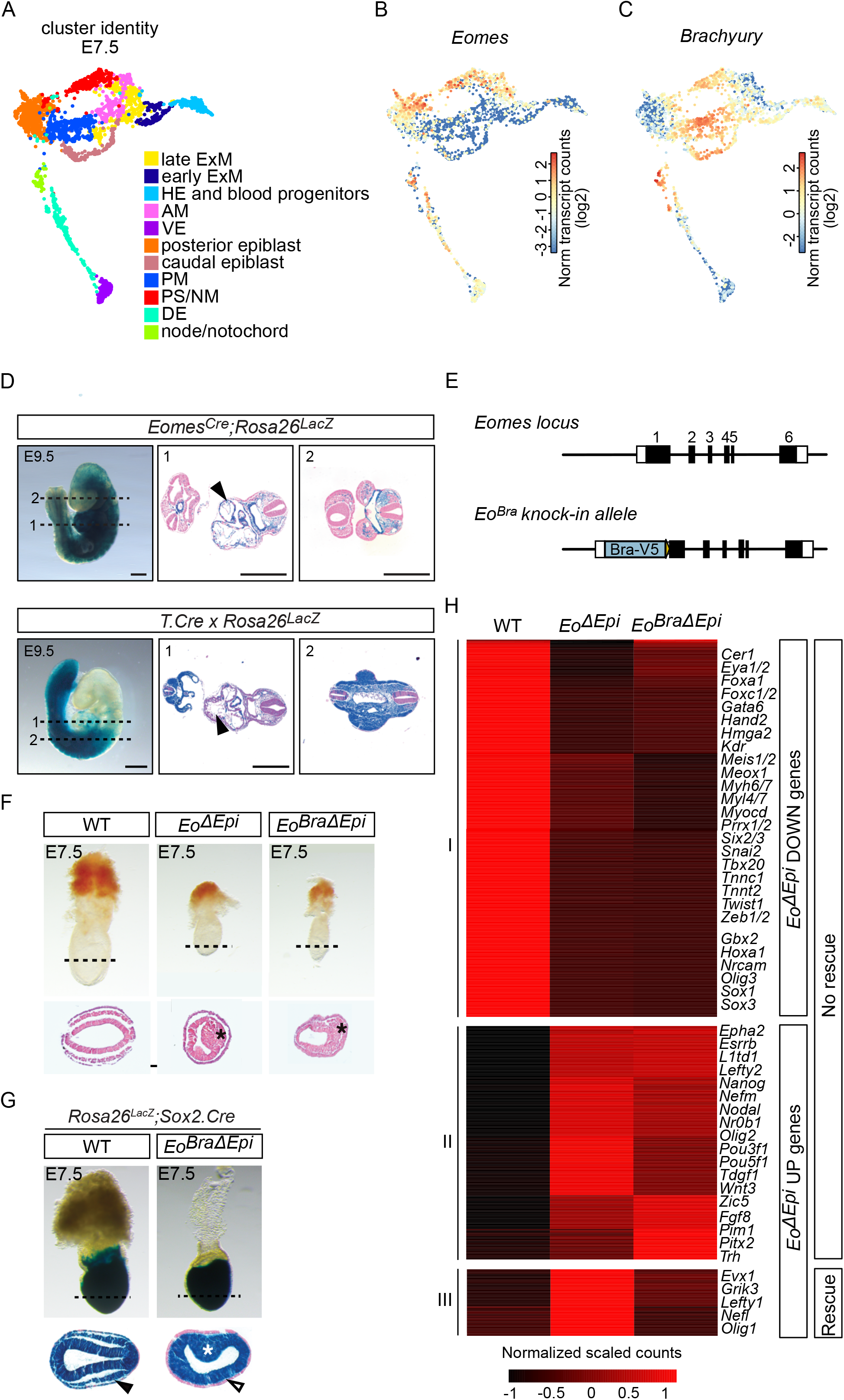
*Eomes* and *Brachyury* specify distinct lineages at different gastrulation stages. **A-C)** UMAPs of scRNA-seq expression analyses of all *Eomes* and/or *Brachyury* expressing cells of embryonic day 7.5 (E7.5) embryos, data from ^28^. (A) Clusters are assigned to indicated cell identities, and (B) *Eomes*, and (C) *Brachyury* expression plotted. Scale bars represent log2 normalized transcript counts. Extraembryonic ectoderm was excluded from UMAPs. ExM extraembryonic mesoderm, HE – haematoendothelium, AM – anterior mesoderm, VE – visceral endoderm, PM – posterior mesoderm, PS/NM – primitive streak/nascent mesoderm, DE – definitive endoderm. **D)** Genetic lineage tracing of *Eomes* and *Brachyury* expressing cells by *Eomes*^Cre^;*Rosa26*^LacZ^ and *T*.*Cre*;Rosa26^LacZ^, respectively. *Eomes*^Cre^;*Rosa26*^LacZ^ fate-labelled cells (upper panel) are stained in blue by β-galactosidase and are found in whole mounts and sections at indicated levels of E9.5 embryos predominantly in anterior embryonic regions, cranial and cardiac mesoderm, vasculature and the gut tube. Descendant cells of *T*.*Cre*;Rosa26^LacZ^ *Brachyury*-expressing cells (lower panel) are found mostly in the posterior portion of the embryo. Scale bars 500 μm. Dashed lines indicate level of transverse sections, black arrowheads highlight differences in staining of cardiac mesenchyme between *Eomes*^Cre^ and *T*.Cre labelled cells. **E)** Schematic of the targeting strategy to insert *Brachyury*-*V5* (*Bra-V5*) coding sequence into the first exon of the *Eomes* genomic locus. Numerics indicate exons. **F)** Whole mount embryos and hematoxylin and eosin stained transverse sections of *Eomes*^ΔEpi^ (*Eomes*^CA/-^;*Sox2*.Cre) and *Eomes*^BraΔEpi^ E7.5 embryos show similar morphogenetic defects, such as failure to form the mesoderm layer and mesoderm cells accumulate in the primitive streak (asterisks). **G)** Fate-labelling using *Rosa26*^*LacZ*^*;Sox2*.*Cre* indicates blue, β-galactosidase-stained epiblast-derived cells at E7.5. *Eomes*^BraΔEpi^ embryos remain surrounded by visceral endoderm (VE, empty arrowhead), due to the failure to form definitive endoderm (DE) cells. The endoderm layer of WT embryos is composed of epiblast-derived DE cells (black arrowhead). **H)** Heatmap showing clustered, differentially expressed genes (adjusted p-value<0.05; log2(FC)>1.0 for upregulated genes, log2(FC)<-1.0 for downregulated genes) between WT, *Eomes*^Δepi^ (*Eo*^Δepi^) and *Eomes*^BraΔEpi^ (*Eo*^BraΔEpi^) embryos analyzed at E7.5 by RNA-seq. Scale represents centered scaled counts normalized by library size and gene-wise dispersion. Key developmental genes are indicated for each cluster.

To compare the tissue contribution of *Eomes-* and *Brachyury*-expressing cells, we performed genetic lineage-tracing by crossing either *Eomes*^Cre^, or *T*.Cre mice to the Cre-inducible *Rosa26*^LacZ^ reporter line and analyzed X-Gal staining at E9.5 to follow the progeny of *Eomes*-or *Brachyury*-expressing cells. *Eomes*-expressing cells give rise to the head mesenchyme, heart and vasculature, and the entire gut tube (Figure 1D) as previously reported ^6^. In contrast, *T*.Cre labelled cells are predominantly found in the posterior portions of the embryo ^30^, including the limb buds and tail region, consistent with functional requirements of *Brachyury* for mesoderm formation posterior to the 7^th^ somite as evident from *Brachyury* homozygous mutant embryos ^12,15^. Anterior mesoderm derivatives including the heart and the cranial mesenchyme are only incompletely labelled in *T*.Cre, *Rosa26*^LacZ^ reporter embryos (Figure 1D, arrowheads), contrasting previous studies that suggested a functional contribution of *Brachyury* to the specification of cardiogenic mesoderm ^31,32^ similar to described *Eomes* functions ^6^. Thus, we re-investigated early cardiac marker gene expression of *Mesp1* and *Mlc2v (Myl2)* in *T*^2J/2J^ *Brachyury* mutant embryos by two-color whole mount *in situ* hybridization in E7.0 to E8.5 embryos (Figure S1J). Until E7.5 *T*^2J/2J^ mutant embryos show little morphological alterations but could be identified by the absence of *Brachyury* expression (Figure S1J, arrowheads). At E7.0 and E7.5 the expression of *Mesp1* in the mid to distal, cardiogenic PS region are similar between WT and *T*^2J/2J^ mutant embryos, while *Mesp1* expression in the proximal PS is reduced (Figure S1J, empty arrowheads), indicating the region giving rise to the extraembryonic mesoderm. Similarly, at E8.5 the expression of *Mlc2v (Myl2)* marking early cardiomyocytes shows little differences in expression levels in the forming heart between WT and *T*^2J/2J^ mutant embryos (Figure S1J). These data indicate that *Brachyury* is dispensable for cardiac specification in line with previously described *Brachyury* mutant phenotypes, where heart morphogenesis is altered secondary to embryonic midline defects, while cardiac lineage specification is undisturbed ^33^.

Next, to compare functions of EOMES and BRACHYURY in global cell specification programs we analyzed the transcriptional profiles of *Eomes*^ΔEpi^, *T*^2J/2J^ and WT embryos from bulk RNA-seq (Figures S2A and B). At E7.5 the deletion of *Eomes* had a marked effect on global gene expression, resulting in 260 downregulated and 147 upregulated genes compared to WT embryos (Figure S2A). In particular, cardiac mesoderm markers (*Myocd, Myl7, Myh7, Tnnt2*), mesoderm markers (*Foxc2, Meis1, Hand2*), and endoderm genes (*Cer1, Cxcr4, Gata6, Foxa1*) are severely downregulated in *Eomes* mutants. Surprisingly, at E7.5 *T*^2J/2J^ mutants show only minor changes in global gene expression compared to WT (Figure S2B), suggesting that *Brachyury* functions are mostly dispensable during the first day of gastrulation (E6.5-E7.5). In contrast, expression data of E8.5 *Brachyury* deficient embryos from published RNA-seq datasets revealed 735 downregulated and 879 upregulated genes of which 70 and 53 were chromatin-bound by BRACHYURY, respectively ^34^. Taken together, these results further emphasize that *Eomes* regulates gene expression during onset of gastrulation (E6.5-E7.5), while *Brachyury* fulfills functions after E7.5 for the generation of posterior mesoderm derivatives.

We next tested whether functional differences between *Eomes* and *Brachyury* result from different expression dynamics, such as earlier and more widespread expression of *Eomes* (Figures S1D-F). Thus, to investigate whether *Brachyury* can compensate for *Eomes* in the epiblast when expressed from the same locus, we generated a targeted knock-in allele of the *Brachyury* coding sequence into the first exon of the *Eomes* genomic locus (*Eomes*^Bra^; Figure S2C). Faithful expression of *Brachyury* from the *Eomes* locus was confirmed (Figures S2D and E), and the *Eomes*^Bra^ allele was intercrossed with the Sox2:Cre deleter strain together with the conditional *Eomes*^CA^ allele to generate *Eomes*^BraΔEpi^ embryos ^5,35^. Here, *Brachyury* is expressed from one *Eomes* allele while the other allele is conditionally deleted from the epiblast (Figure 1E and S2C). *Eomes*^BraΔEpi^ embryos showed similar morphogenetic defects as found in *Eomes*^ΔEpi^ embryos (Figure 1F), such as cells that fail to migrate from the primitive streak (PS) to form the mesoderm wings and instead accumulate in the region of the mutant PS ^5^. Also, marker analysis by *in situ* hybridization indicated that *Brachyury* expression from the *Eomes* locus fails to rescue the cell specification defects found in *Eomes*^ΔEpi^ embryos resulting in the absence and/or severe reduction of markers for cardiac mesoderm (*Mesp1, Myl7*) and DE (*Foxa2*), while PS markers *Wnt3a* and *Fgf8* remain strongly expressed in *Eomes*^BraΔEpi^ embryos (Figure S2F). To test whether epiblast derived cells contribute to the DE cell lineage in *Eomes*^BraΔEpi^ embryos, we additionally genetically fate labelled epiblast cells by intercrossing the *Rosa26*^*LacZ*^ reporter and performed β-Galactosidase staining to detect Sox2:Cre-labelled epiblast-derived cells (Figure 1G). In WT embryos the majority of DE cells in the endoderm layer are labelled at E7.5, indicating their origin from the epiblast (Figure 1G; ^36^). In contrast, in *Eomes*^BraΔEpi^ embryos the endoderm cell layer is composed of unlabeled VE cells that are not replaced by DE cells (Figure 1G), which is also reflected by persisting expression of the VE marker *Afp* overlaying the epiblast (Figure S2F).

The inability of *Brachyury* to compensate for the functional loss of *Eomes* in the epiblast is further confirmed by transcriptional profiles by bulk RNA-seq of E7.5 *Eomes*^BraΔEpi^ and *Eomes*^ΔEpi^ embryos compared to WT (Figure 1H, Table S1). Clustering analysis of differentially expressed genes identified three major groups (Figure 1H): Cluster I consists of a large group of 326 genes downregulated in *Eomes*^ΔEpi^ compared to WT embryos that are not rescued in *Eomes*^BraΔEpi^ embryos, including early key AM and DE markers (Figure 1H, Cluster I). Clusters II and III contain genes with increased expression in *Eomes*^ΔEpi^ compared to WT embryos. Cluster II genes are not repressed by prematurely expressed *Brachyury* in *Eomes*^BraΔEpi^ embryos and include markers for anterior epiblast (*L1td1, Nr0b, Epha2*) and early NE (*Pou3f1, Olig2*) (cluster II, 200 genes). Relatively fewer genes with increased expression in *Eomes*^ΔEpi^ were rescued in *Eomes*^BraΔEpi^ embryos and are represented in cluster III (57 genes). Among those were anterior epiblast (*Lefty1* and *Evx1*) and NE marker genes (*Girk3, Nefl* and *Olig1*), indicating a partial redundancy between *Eomes* and *Brachyury* to repress pluripotency and NE genes in the early epiblast as previously described ^21^. In summary, despite partially overlapping expression in the ME-forming epiblast in early gastrulating embryo, *Eomes* and *Brachyury* fulfill different functions during lineage specification and morphogenesis, and *Brachyury* does not compensate for loss of *Eomes* functions even when expressed under the same genomic control *in situ*.

### Sets of specific *Eomes* and *Brachyury* target genes and programs

To define a high-confidence set of *Eomes*- and/or *Brachyury*-dependent target genes that confer lineage specific programs during ME differentiation we took advantage of a tightly regulatable *in vitro* embryoid body (EB) differentiation system (Figure 2A). EBs were formed from mouse embryonic stem cells (mESCs) with single (sKO, EoKO or BraKO) or combined genetic deletions of both *Eomes* and *Brachyury* (dKO)(Figure 2B). To fully control Tbx TF expression independent of upstream signals, we reintroduced dox-inducible expression constructs for GFP-tagged cDNAs (EoGFP and BraGFP)(Figure 2B) that phenotypically and molecularly rescue the genetic deletions ^21^. ESC aggregates were formed for 48 hours in basal medium, resembling the priming phase of pluripotency, before differentiation was induced with the NODAL-analogue ActivinA (ActA) and -/+ Doxycyclin (Dox) to induce the expression of EoGFP/BraGFP in sKO or dKO cells (Figures 2A and B).

**Figure 2.**
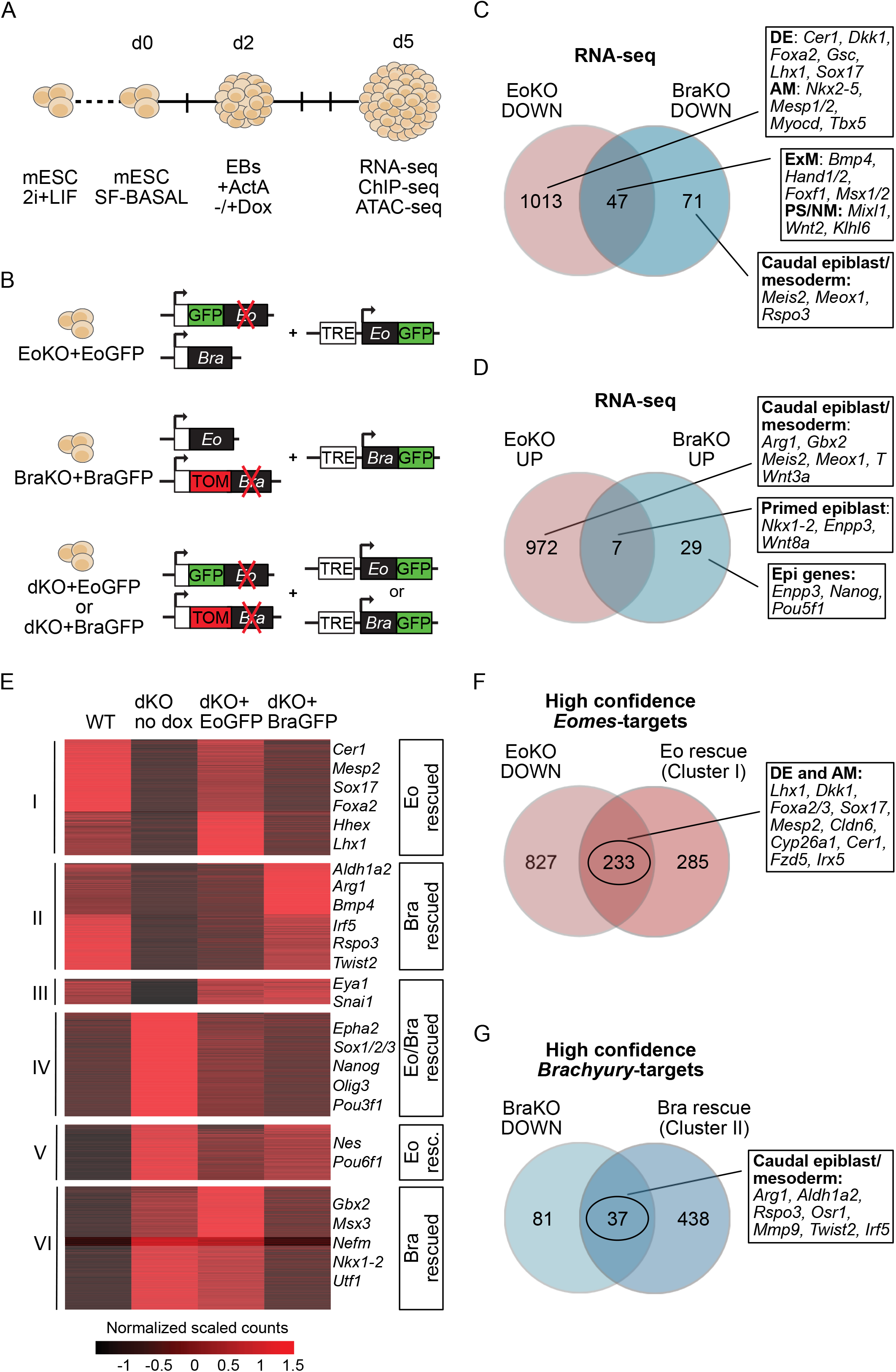
Sets of specific *Eomes* and *Brachyury* target genes and programs. **A)** Schematic illustration of embryoid body (EB) differentiation. mESC – mouse embryonic stem cells; SF – serum free; ActA – ActivinA; d – day of differentiation; Dox – doxycycline. **B)** Schematic illustration of different cell lines used: Cells with single gene deletions of *Eomes* (EoKO) or *Brachyury* (BraKO), and double KO (dKO) cells were generated by the insertion of fluorescent reporters on one allele (*Eomes*^GFP^ and *Brachyury*^Tomato^) and an out-of-frame deletion on the second allele for each gene. EOMES-GFP or BRACHYURY-GFP were introduced into the doxycycline-inducible locus (TRE) of EoKO, BraKO or dKO cells (EoKO+EoGFP, BraKO+BraGFP, dKO+EoGFP, dKO+BraGFP). **C, D)** Venn diagram of RNA-seq analyses showing the overlap of downregulated (C) or upregulated (D) genes in EoKO and in BraKO compared to WT EBs (adjusted p-value<0.05, log2(FC)>1.5 for upregulated genes, log2(FC)<-1.5 for downregulated genes). Key markers for different cell lineages are indicated on the right. DE – definitive endoderm, AM – anterior mesoderm, ExM – extraembryonic mesoderm, PS/NM – primitive streak/nascent mesoderm, Epi – epiblast. **E)** Heatmap showing normalized scaled counts of clustered, differentially expressed genes between WT and dKO cells (no dox), and rescue of expression levels by induction of EoGFP (clusters I and V) or BraGFP expression (clusters II and VI), or both (cluster III and IV). Adjusted p-value<0.05; log2(FC)>1.0 for upregulated genes, log2(FC)<-1.0 for downregulated genes. Scale represents centered scaled counts normalized by library size and gene-wise dispersion. Relevant marker genes in each group are indicated. **F, G)** High confidence target genes of *Eomes* or *Brachyury* defined by intersecting genes that are downregulated in EoKO (F) or BraKO (G) with EoGFP-rescued and BraGFP-rescued genes (E), respectively. Key markers for different cell lineages are indicated. DE – definitive endoderm, AM – anterior mesoderm.

First, we compared gene expression in EoKO and BraKO single mutant EBs by RNA-seq. Similar to the transcriptional profiles of E7.5 embryos (Figures S2A and B) EoKO EBs showed a higher number of differentially regulated genes than BraKO EBs, i.e. 1,060 downregulated and 979 upregulated genes in EoKO, and only 118 downregulated and 36 upregulated genes in BraKO compared to WT, respectively (Table S2; Figures 2C and D). Only a small portion of deregulated genes in EoKO were also deregulated in BraKO, indicating small overlap in the gene regulatory programs specified by both Tbx factors in these differentiation conditions (Figures 2C and D; Table S2). Among downregulated genes in EoKO compared to WT were regulators of cardiac cell differentiation (*Nkx2-5, Gata4, Myocd, Hand1, Tbx5*), endoderm (*Cer1, Foxa2, Hhex, Sox17)*, and early regulators of gastrulation (*Nodal, Tdgf1, Mixl1*) (Figure 2C and Table S2). Downregulated genes in BraKO EBs compared to WT EBs were markers for paraxial mesoderm and caudal epiblast/tail bud (*Rspo3, Meox1*), and markers that are shared between cardiac and extraembryonic mesoderm (*Gata5, Hand1/2, Msx1*) (Figure 2C and Table S2). Interestingly, *Brachyury* (*T*) and markers of the posterior mesoderm gene program (*Arg1, Meox1, Meis2, Wnt3a*) showed increased expression in EoKO cells compared to WT (Figures 2D and S3A; Table S2), suggesting that co-expression with EOMES restricts BRACHYURY functions. In contrast, expression of *Eomes* and its target genes was not increased in BraKO EBs (Figures 2D and S3B; Table S2) suggesting that repression of *Eomes* and *Brachyury* is not mutual and that repression by *Eomes* serves to inhibit premature activation of *Brachyury*-induced programs.

Next, we profiled gene expression of EBs containing dox-inducible expression constructs of GFP-fused *Eomes* or *Brachyury* cDNAs (dKO+EoGFP and dKO+BraGFP) in the dKO genetic background (Figure 2B). In the absence of Tbx expression dKO EBs entirely lack the potency to specify any ME lineages, retain pluripotency and eventually differentiate towards NE cell types ^21^. Following induced expression most of the differentially expressed genes in dKO (no dox) cells compared to WT were rescued after either *Eomes* (clusters I and V) or *Brachyury* induction (clusters II and VI)(Figure 2E, and Table S3). However, only a small number of downregulated genes in dKO EBs were rescued by both TFs (Figure 2E, n=113, cluster III). Interestingly, a relatively high proportion of upregulated genes in dKO (no dox) (n= 462, cluster IV) was repressed upon both *Eomes* and *Brachyury* re-expression, indicating the relatively higher level of redundancy to repress NE and pluripotency genes once ME gene programs are induced by Tbx and other ME TFs ^21^.

The majority of cardiac mesoderm and DE genes were rescued by EoGFP re-expression (cluster I, Figure 2E), while BraGFP induced expression of paraxial mesoderm, and skeletal muscle markers (cluster II, Figure 2E). Gene ontology (GO) analysis of downregulated genes in EoKO (FigureS3C) and BraKO EBs (FigureS3D) reflected this observation showing that distinct gene programs are initiated early during ME lineage specification by either EOMES or BRACHYURY, contrasting previous views of the primary formation of a still unspecified common ME progenitor. To define a high-confidence set of specific target genes of *Eomes* and *Brachyury* we intersected downregulated genes in sKO EBs (EoKO or BraKO) with genes that were rescued in dKO EBs by the induced expression of *Eomes* or *Brachyury*, respectively (Figures 2F and G). These high-confidence targets recapitulate the suggested and observed regulation of AM and DE gene programs by *Eomes*, and posterior mesoderm derivatives by *Brachyury* (Figures 2F and G). In summary, the use of an EB-based, tightly controllable differentiation model allowed to dissect the target genes that guide distinct, lineage specific genetic programs regulated by *Eomes* and *Brachyury*.

### EOMES and BRACHYURY show complex and largely non-overlapping binding patterns at target genes

Previous studies suggested overlapping genomic binding of Tbx factors to similar consensus binding motifs ^18,23^. To refine sets of EOMES and BRACHYURY specific, directly regulated target genes, we analyzed DNA occupancy by chromatin immunoprecipitation coupled with sequencing (ChIP-seq) of day 5 differentiated EBs using EoKO+EoGFP and BraKO+BraGFP ESCs (Figure 2B)^21^. Here, ChIP with the same antibody directed against the GFP-tag in controlled expression conditions allows for the comparative analysis of chromatin binding of both Tbx factors in identical differentiation conditions (Figures 3A and S4A). The genome-wide analysis identified a high number of common sites co-bound by EOMES and BRACHYURY (8,601 regions), and also 10,932 EOMES (EO)-unique and 2,091 BRACHYURY (BRA)-unique binding sites (Figures 3A, B, and S4B). To analyze EOMES or BRACHYURY directly, cis-regulated genes, we used GREAT (Genomic Regions Enrichment of Annotations Tool, ^37^) to associate genes with EOMES- and BRACHYURY-bound regions (Figure S4C, Table S4). The interrogation of binding-site associated genes revealed complex patterns of EOMES and BRACHYURY binding to gene regulatory sites, so that target genes often contained multiple bound regions, that can be common or unique for each Tbx factor (Figures 3A, C, and S4A), and different combinations of Tbx-bound regions are found among target genes (Figure 3C, Table S5). Thus, despite the large number of EO- and BRA-unique binding sites, the associated target genes of EOMES-or BRACHYURY-bound sites grossly overlapped (Figure S4C, Table S4). However, irrespective of the extensive overlap in ChIP-binding associated target genes, we found a striking correlation of unique binding sites with specifically EOMES or BRACHYURY-regulated genes (Figure 3C). For example, *Msgn1* and *Sox18* possess BRA-unique, and *Hhex1, Mesp1* and *Nkx2-1* harbor EO-unique binding sites (Figures 3A and S4A) and are specifically regulated by either TF, respectively. These complex binding patterns of EOMES and BRACHYURY suggest that they regulate large common sets of genes, but at the same time govern fine-tuning of precise spatio-temporal expression patterns within distinct gene programs. Next, we wanted to confirm that unique binding of EOMES or BRACHURY to regulatory sites (Figure 3B) indeed correlates with expression of associated genes. Thus, we compared the groups of associated genes that harbor different combinations of EOMES and BRACHYURY binding (Figure 3C) with the lists of high-confidence *Eomes*-or *Brachyury*-target genes (Figures 2F and G). Therefore, we plotted the frequency of how often genes associated to different binding patterns are also specifically regulated by *Eomes* or *Brachyury* (Figure S4D). This analysis shows that about 45% of genes associated to EO-unique peaks only also belong to the high-confidence group of *Eomes* target genes, while only about 10% of genes with only common peaks fall into the group of high-confidence *Eomes* targets. Similarly, BRA-bound genes (Common+Bra-uni and Bra-uni only group) are more frequently found in the group of high confidence *Brachyury* than high-confidence *Eomes* target genes (Figure S4D), however, at much reduced frequencies under used experimental conditions of ActivinA stimulation that revealed only relatively few *Brachyury* target genes (Figure 2G).

**Figure 3.**
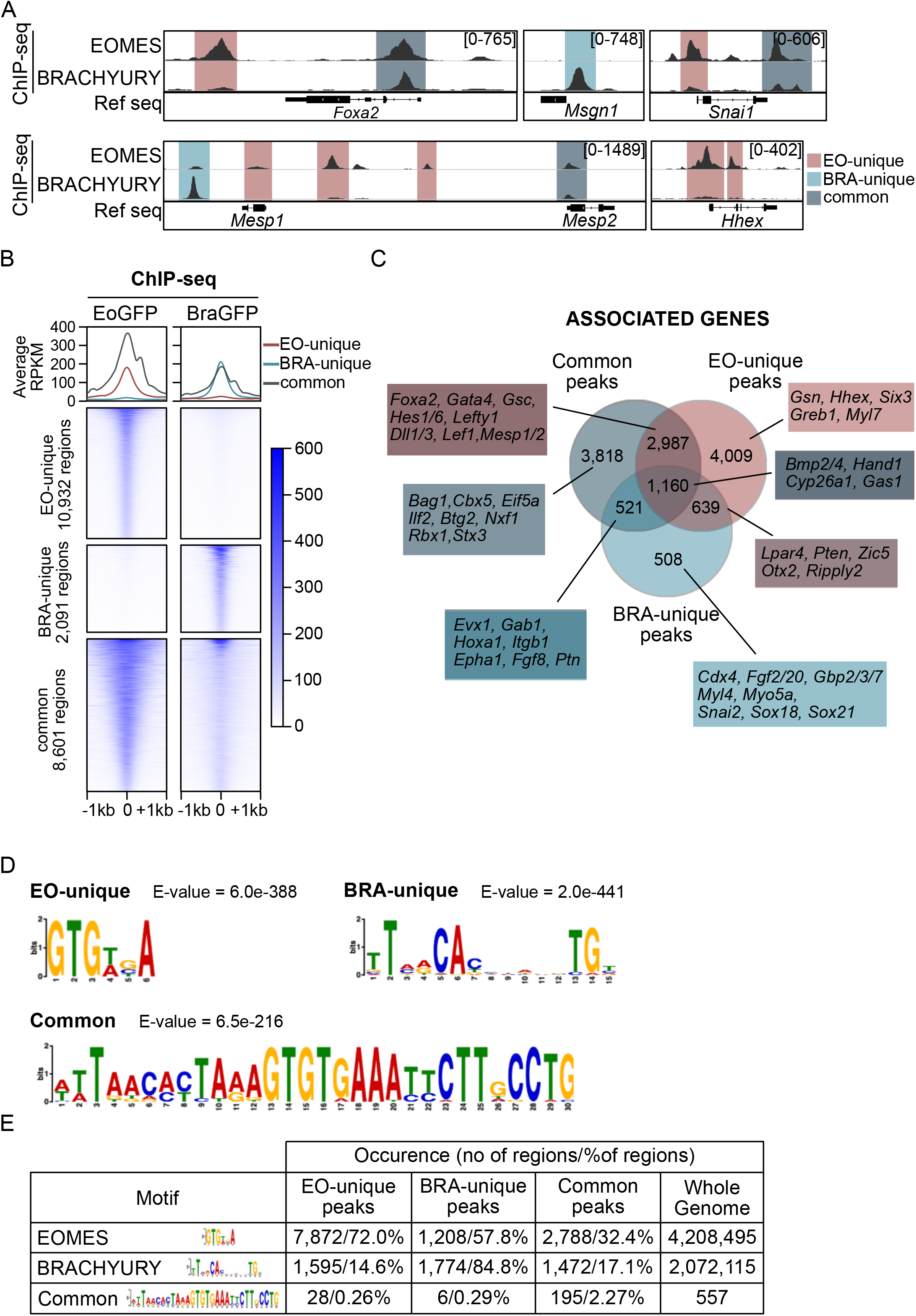
EOMES and BRACHYURY show complex and largely non-overlapping binding patterns at target genes. **A)** Examples of ChIP-seq coverage tracks of EOMES- 9and/or BRACHYURY-bound region close to target genes (*Foxa2, Msgn1, Snai1, Mesp1/2, Hhex*) in ActvinA-induced EBs (EoKO+EoGFP and BraKO+BraGFP). Counts normalized to RPKM are indicated. **B)** Heatmaps of specifically EOMES (EO-unique), BRACHYURY (BRA-unique) and commonly bound EOMES+BRACHYURY (common) binding regions from ChIP-seq in EoKO+EoGFP and BraKO+BraGFp EBs. Scale represents normalized counts (RPKM) for ChIP-seq ±1 kb centered around the peak. **C)** Venn diagram showing the intersection of associated genes of common, EO- and BRA-unique ChIP-binding peaks. Examples of associated genes are indicated for each type of peak distribution. **D)** Position weight matrices of sequence motifs for Tbx factor binding generated by *de novo* motif prediction in EO-unique, BRA-unique and common binding peaks. E-values are indicated. **E)** Absolute numbers of occurrence and relative presence of EOMES, BRACHYURY and common binding motifs in unique and common ChIP-peaks compared to appearance in the genome. Percentage of indicated region is calculated by the number of motifs in relation to the total number of unique/common regions.

Next, we investigated what determines binding specificity of EOMES and BRACHYURY to uniquely bound sites by analyzing the underlying Tbx consensus binding motifs. We performed *de novo* motif analysis of the EO-unique, BRA-unique and EO+BRA common peaks using MEME suite ^38^, which revealed distinct motifs with different lengths and base compositions (Figure 3D) between EO-unique, BRA-unique and common bound regions. EO-unique regions contained an enriched motif of 6 bases, including 6 of the 7 bases of the known Tbx binding sequence GGTGTGA ^39^. The motif enriched in BRA-unique peaks resembled the previously described palindromic motif of 16 bases ^22^. The analysis of common peaks revealed a remarkably long sequence motif of 30 bases containing both motifs for EOMES and BRACHYURY, that are also found within the unique peaks (Figure 3D). As expected, predicted motifs are frequently present in ChIP-peaks so that the EOMES motif is found in 72% of all EO-unique peaks, and the BRACHYURY motif in 85% of BRA-unique peaks (Figure 3E). Interestingly, while the EOMES motif was found in 58% of BRA-bound peaks, the BRACHYURY motif is only found in 15% of the EO-bound peaks suggesting that EOMES might potentially impact on *Brachyury*-targets, but not in reverse. Therefore, EOMES might bind to BRA-unique sites to compensate for BRACHYURY function in absence of BRACHYURY. However, in absence of EOMES, BRACHYURY cannot bind to EOMES-sites and compensate for EOMES function as seen in *Eomes*^BraEpi^ embryos (Figures 1F-H and S2F). Of note, while all identified motifs show the basic sequence described for T-box DNA binding domains ^18,23^, the specific motifs found in EO- and BRA-unique binding regions are not necessarily overlapping in sequence and could provide the basis to explain EOMES and BRACHYURY-binding specificity to their regulatory sites. Alternatively, it was suggested that Tbx binding specificity could be regulated by co-operative binding with distinct other TFs ^19^. Thus, we analyzed annotated TF binding motifs enriched in EO-unique over BRA-unique binding and *vice versa* (HOMER ^40^, Figures S4E, F). EO-unique regions showed an enrichment of motifs for FOXA1/2, GATA1/2/4/6, FOXD3 and TBX6, which are known regulators of ME development, and a TBOX:SMAD binding motif reflecting cooperative regulation of target genes by EOMES and SMAD signaling ^8^. In contrast, motifs of members of the WNT and JNK signaling pathways were found in BRA-unique regions suggesting different signaling co-factors that co-operatively function together with EOMES and BRACHYURY. However, since ChIP binding profiles were measured in identical EB differentiation conditions, it is unlikely that signal-induced factors, such as SMADs or TCF/LEF factors impacts on Tbx binding specificity between EOMES and BRACHYURY, but more likely play a role in subsequent steps of transcriptional activation. In summary, the analysis of binding specificities under fully controlled and comparable experimental conditions suggests that the binding specificity of EOMES and BRACHYURY is indeed based on different affinities to specific DNA consensus sites of Tbx-specific gene-regulatory elements with EOMES showing the potential to bind to BRACHYURY target sites.

### EOMES and BRACHYURY unique binding sites locate to enhancers and control specific target gene expression

To examine how overlapping and unique binding sites of EOMES and BRACHYURY contribute to the regulation of target genes, we analyzed the distribution of binding sites in respect to target gene genomic features (Figure 4A). Interestingly, the distribution of common and unique sites significantly differs, so that common binding sites are predominantly located in promoter regions while unique EO- and BRA-bound sites correspond to intronic and intergenic regions, likely acting as distal enhancer elements (Figure 4A). This supports the idea that EOMES and BRACHYURY share common roles in promoter activation but may exert unique functions in tissue- and temporal-specific enhancer regulation of transcription.

**Figure 4.**
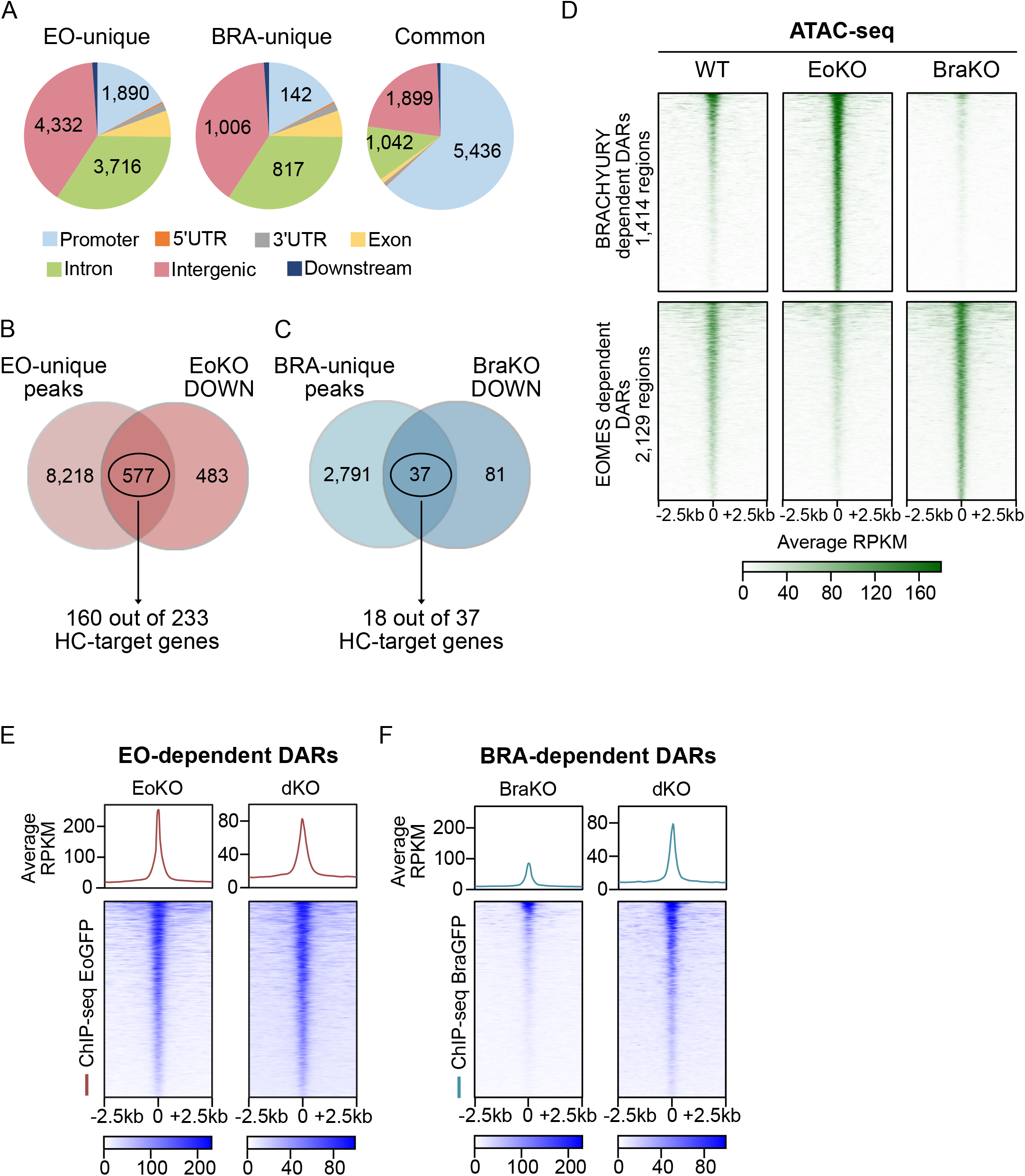
EOMES and BRACHYURY unique binding sites locate to enhancers and control specific target gene expression. **A)** Pie charts showing the distribution of EO-unique, BRA-unique and common ChIP binding-peaks along different genomic regions. Common binding peaks are predominantly found in promoters, unique binding sites show a high prevalence in intronic and distal intergenic regions. Absolute numbers are indicated for the different genomic regions. **B, C)** Venn-diagrams showing the degree of overlap of genes associated to EO-(B) or BRA-unique (C) peaks and genes downregulated in EoKO and BraKO by RNA-seq, respectively. Overlapping genes resemble EOMES or BRACHYURY-regulated genes that are also largely represented in the high confidence sets of target genes as indicated. HC – high confidence. **D)** Heat maps of EOMES- and BRACHYURY-dependent differentially accessible regions (DARs) in WT, EoKO and BraKO EBs showing decreased accessibility of EOMES-dependent DARs in EoKO compared to WT, but increased accessibility in BraKO cells. BRA-dependent DARs are significantly more accessible in EoKO and almost absent in BraKO EBs. Scale represents normalized counts (RPKM) ±2.5 kb around the center of the peak. **E, F)** Comparative ChIP-binding of EoGFP and BraGFP in the presence (EoKO+EoGFP and BraKO+BraGFP (single KO, sKO), or absence of the other Tbx factor (dKO +EoGFP/BraGFP) shows no major changes in EoGFP-binding in different cells, while BraGFP-binding is markedly increased in dKO EBs compared to BraKO cells. Scale represents normalized counts (RPKM) ±2.5 kb around the center of the peak.

To further underscore that unique EOMES and BRACHYURY binding mostly contributes to specific target gene regulation, we intersected the previouRemarkably, of all EOMES-bound sites sly experimentally identified *Eomes* or *Brachyury* regulated genes (DEGs in EoKO and BraKO EBs, Figures 2C and D) with uniquely bound regions (Figures 3B and C). Of 1060 downregulated genes in EoKO cells, 577 had an EO-unique binding peak (Figure 4B). Within this group of genes were 160 of the 233 genes that were identified as high-confidence *Eomes* target genes (Figure 2F). Overlapping BRA-unique peaks with genes downregulated in BraKO cells resulted in 37 genes of which 18 were classified as high confidence *Brachyury* target genes (Figures 4C and 2G). In conclusion, target gene specificity between *Eomes* and *Brachyury* is conferred by unique binding to different enhancer elements to regulate the transcription of the associated genes, and most likely not by shared functions at promoter elements.

### EOMES restricts BRACHYURY functions and competing gene programs

Transcriptional profiles of *Eomes-* and *Brachyury*-expressing cells showed grossly different regulation of alternative cell specification programs (Figure 2E). However, in the E7.0 embryo both Tbx TFs are coexpressed in cells at the PS (Figures S1E and F). Thus, we asked if there is mutual regulation between EOMES and BRACHYURY to discriminate competing cell specification programs. First, we analyzed if EOMES impacts on chromatin binding of BRACHYURY and *vice versa* by comparing chromatin-binding of EOMES in *Bra*-proficient (EoKO+EoGFP) and *Bra*-deficient (dKO+EoGFP) cells and performed the reciprocal analysis for BRA-binding (Figures S5A and B). Remarkably, of all EOMES-bound sites (n= 40,037 peaks) only 34 were differentially bound in EoKO+EoGFP and dKO+EoGFP cells, demonstrating that EOMES-binding is not influenced by the presences of BRACHYURY (Figure S5A). In contrast, BRACHYURY-binding (n= 12,702 peaks) is modulated by the presence of EOMES and we found differential binding at 1,855 sites whereby 1,673 regions show reduced binding in the presence of EOMES (Figure S5B).

Next, we wanted to test if there is also mutual regulation of molecular and gene regulatory functions between EOMES and BRACHYURY. Previous studies suggested that Tbx factors control the establishment of chromatin accessibility at target gene regulatory sites ^21,25,41^. Thus, we used ATAC-seq (Assay for Transposase-Accessible Chromatin with sequencing) as read-out for EOMES and BRACHYURY functions measured by increased chromatin accessibility. We compared differentially accessible chromatin regions (DARs) in EoKO and BraKO differentiated EBs and defined 1,414 BRACHYURY-dependent (sites with reduced chromatin accessibility in BraKO cells) and 2,129 EOMES-dependent (sites with reduced chromatin accessibility in EoKO cells) DARs (Figures 4D and S5C). We plotted ATAC-signals of BRACHYURY- and EOMES-dependent DARs in WT, EoKO and BraKO EBs (Figure 4D). As expected, EOMES-dependent DARs and BRACHYURY-dependent DARs showed reduced accessibility in EoKO and BraKO EBs, respectively (Figure 4D). However, BRACHYURY-dependent DARs showed a strong increase of ATAC-signals in EoKO EBs, while EO-dependent DARs were only slightly increased in BraKO cells (Figure 4D), indicating enhanced BRACHYURY functions in EoKO compared to WT EBs. Next, we analyzed EOMES and BRACHYURY chromatin-binding to EOMES- and BRACHYURY-dependent DARs in the presence (sKO, either EoKO or BraKO) or absence (dKO) of the other Tbx factors (Figure 4E and F). Here, we found that also at BRA-dependent DARs BRACHYURY-binding is inhibited in the presence of EOMES (BraKO+BraGFP cells) when compared to dKO cells (Figure 4F), while the presence of BRACHYURY (EoKO cells) had less effects on EOMES binding when compared to dKO cells (Figure 4E), reflecting the previous genome-wide binding analysis in sKO and dKO cells (Figure S5A, B). In conclusion, these findings demonstrate that EOMES restricts BRACHYURY chromatin binding and chromatin accessibility at BRA-bound sites. In reverse, BRACHYURY does not significantly impact on EOMES chromatin binding, has only minor effects on chromatin accessibility of EOMES-regulated genes, and also the absence of *Brachyury* does not alter transcription of *Eomes*-regulated genes in either EB differentiation experiments (Table S2; Figures 2C and D) or in early embryos preceding E7.5 (Figures S2B). To further corroborate the observation of antagonistic *Eomes*-functions on *Brachyury* activities, we employed another *in vitro* embryoid differentiation model which reflects the early experimental phase of gastruloid formation from mESCs, that strictly relies on *Brachyury* gene functions ^42,43^. Here, EBs are induced by an 24 hours pulse of the small molecule GSK3b-inhibitor CHIR to activate the WNT signalling pathway that induces *Brachyury* expression (Figure S5D ^44,45^). We compared CHIR-induced EBs generated from WT cells with EBs generated from WT+EoGFP cells that were induced to express EoGFP during the CHIR pulse. Both WT and WT+EoGFP EBs showed similar levels of nuclear BRACHYURY (Figure S5E). However, the expression of *Brachyury-*regulated target genes *Aldh1a2, Rspo3, Msgn1* and *Tbx6* was severely reduced upon induced *Eomes*-expression, while the *Eomes* high-confidence, AM target gene *Cyp26a1* (Figure 2F) showed increased expression levels (Figure 5A), supporting the view that EOMES restricts BRACHYURY functions. In line with these findings, also BRACHYURY binding to regulatory sites of its target genes is increased in the absence of *Eomes* (dKO+BraGFP) compared to *Eomes*-proficient cells (BraKO+BraGFP; Figure 5B and S5F). In comparison, the *Eomes*-target gene *Cyp26a1* shows a unique binding site for EOMES with reduced chromatin accessibility in EoKO and no change in presence or absence of *Brachyury* (compare to dKO; Figures 5B).

**Figure 5.**
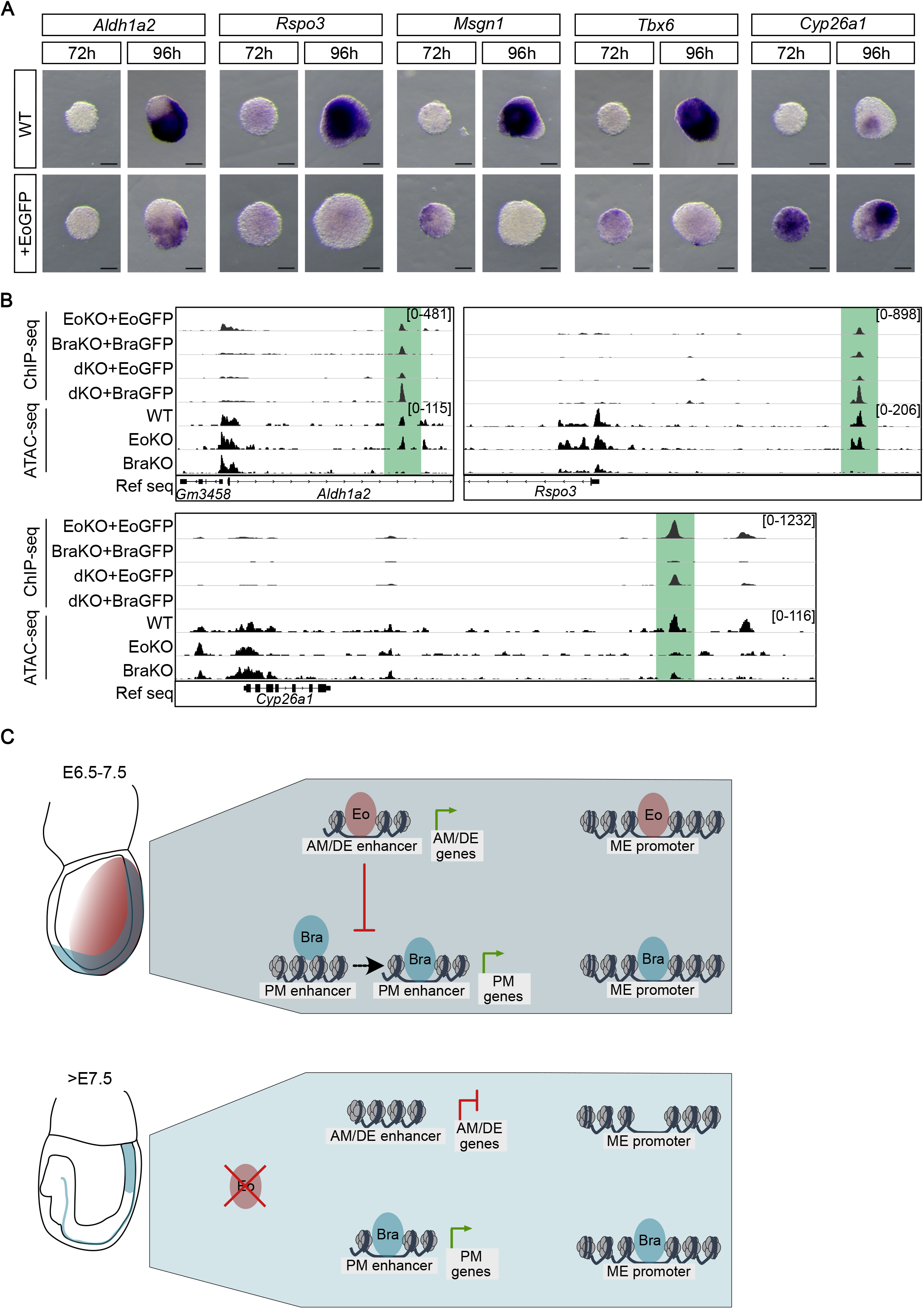
EOMES restricts BRACHYURY functions and competing gene programs. **A)** Whole mount *in situ* hybridization of WT and WT+EoGFP CHIR-induced EBs at indicated timepoints for *Brachyury*-targets *Aldh1a2, Rspo3, Msgn1, Tbx6* and the *Eomes*-target *Cyp26a1* showing that *Brachyury*-target gene expression is reduced upon overexpression of *Eomes*. Scale bars 100 μm, n≥6. **B)** Examples of ChIP-seq coverage tracks of EOMES- and/or BRACHYURY-bound regions in ActvinA-induced EBs (EoKO+EoGFP and BraKO+BraGFP or dKO+EoGFP and dKO+BraGFP) close to *Brachyury*-target genes (*Rspo3* and *Aldh1a2*) or *Eomes*-target gene *Cyp26a1. Brachyury*-target genes show preferential BRACHYURY binding in absence of EOMES and loss of open chromatin in BraKO compared to EoKO and WT as shown by ATAC-seq coverage tracks of WT, EoKO and BraKO ActivinA-induced-EBs. Counts normalized to RPKM are indicated. **C)** Model for lineage-specific activities of EOMES and BRACHYURY. EOMES and BRACHYURY bind in overlapping fashion at promoter regions of common mesoderm and endoderm target genes, while enhancer binding is specific and conferred by different Tbx binding motifs for EOMES and BRACHYURY. *Eomes* activates gene programs that act early during gastrulation to specify AM and DE lineage programs (E6.5-7.5), concomitantly repressing *Brachyury* binding and activities, indicated by reduced enhancer accessibility. After E7.5 *Eomes* expression rapidly declines and the repression of *Brachyury* functions is released, allowing BRACHYURY to instruct lineage specific programs for PM and posterior PS derivatives.

In summary, this study leads to a model of diverging and partially competing functions of the two related Tbx TFs *Eomes* and *Brachyury* to instruct successive gene programs during cell lineage specification at gastrulation (Figure 5C). While target gene sets and binding sites of both TFs are overlapping at promoters, enhancers of target genes are generally specific for one or the other Tbx TF. Hence, *Eomes* activates gene programs that act early during gastrulation to specify AM and DE lineage programs, concomitantly repressing *Brachyury* activities. *Brachyury* functions only become prominent in the following stage of gastrulation, once *Eomes* expression ceases to release repressive activities on *Brachyury*, that acts to specify PM and other derivatives of the PS by instructing different gene programs.

## Discussion

In this study we addressed the transcriptional control of cell specification to mesoderm and definitive endoderm lineages during early stages of mammalian gastrulation. We aimed for defining the gene-expression controlling functions of the Tbx TFs *Eomes* and *Brachyury*, two main regulators of lineage specification and morphogenesis, to resolve a more than two decades old conundrum about specific and overlapping functions of these transcriptional regulators ^19,23,46^. We first delineated the different sets of target genes, that characterize the gene regulatory programs for AM and DE cell lineages that are instructed by *Eomes*, and those for posterior mesoderm lineages induced by *Brachyury*. We previously demonstrated that combined activities of both TFs account for lineage specification to all ME cell types, so that their compound genetic deletion results in the complete absence of any ME, accompanied by gross chromatin inaccessibility of enhancers for ME genes ^21^. While target genes of *Eomes* and *Brachyury* are partially overlapping (this study and Ref. 18-20), we demonstrate that *Brachyury* cannot compensate for the loss of *Eomes* in genetic experiments, showing that functional specificity resides within protein functions and is not due to differences in expression timing or expression domains (Figure 1).

To precisely compare and discriminate molecular functions of both Tbx proteins we generated *Eomes* and *Brachyury* deficient (sKO and dKO) ES cells and re-introduced inducible expression of tagged EOMES and BRACHYURY, faithfully rescuing endogenous gene function ^21^. Hence, we could induce expression of both TFs independent of the upstream signals, e. g. TGFβ (NODAL) or WNT that might impact in a Tbx-dependent and/or - independent fashion on chromatin binding and/or transcriptional responses as previously suggested ^20^. This approach allowed for direct comparison of EOMES and BRACHYURY chromatin-binding patterns and changes in chromatin accessibility as higher order functional readout for target gene regulation. Despite the high degree of amino acid conservation within the T-box DNA-binding domain, the chromatin binding patterns of EOMES and BRACHYURY considerably differ on the genome-wide scale. Common binding of EOMES and BRACHYURY is predominantly seen at promoter-near regions, while binding to promoter-distant intronic/intergenic enhancer sites is largely non-overlapping. Binding to unique enhancer sites confers specificity for target gene regulation, while commonly bound promoter regions are only weakly associated with specificity for target gene expressions as previously shown in mouse gastrulation embryos ^47^. *De novo* TF-binding motif analysis of uniquely bound sites revealed that chromatin binding specificity is achieved through different binding motifs for EOMES and BRACHYURY, reflecting the classical view that TF specificity is ensured by binding to different, sequence specific motifs. Additionally, cooperative binding with other TFs as co-factors could account for specificity as suggested for BRACHYURY and BMP-mediating SMAD1 ^27,48^. TF consensus motif analysis of EOMES and BRACHYURY bound regions indeed showed an enrichment for motifs of AM- and DE-associated TFs in EOMES ChIP-peaks (FOXA1, GATA4, TBOX:SMAD) and Wnt- and mesoderm-associated TFs (TCF3, JUN-AP1, LEF1) close to BRACHYURY-bound sites. This bias in cooperating TFs might additionally contribute to the correct spatio-temporal activation of different lineage programs, such as early specification of AM and DE by Nodal-induced EOMES in the anterior PS ^8,49^, followed by the activation of PM programs by BRACHYURY that is induced and acts together with canonical Wnt-signals in positive feed-forward loops ^44,45,50^.

Although *Eomes* and *Brachyury* are co-expressed in cells of the primitive streak at early stages of gastrulation, embryonic functions of *Eomes* precede those of *Brachyury*, and the genetic deletion of *Brachyury* does not result in obvious phenotypic and/or molecular (as judged by RNA profiling) defects during the first day of gastrulation (E6.5 – E 7.5). Our analyses suggest that the activities of BRACHYURY to regulate target gene expression for posterior mesoderm specification is antagonized by EOMES in coexpressing cells, and we find increased chromatin binding and chromatin accessibility of BRACHYURY-bound regions when *Eomes* is genetically deleted (Figures 4D and F). Moreover, forced expression of EOMES antagonizes the expression of *Brachyury*-targets despite unaltered *Brachyury* expression (Figures 5A and S5E). In reverse, repressive effects of BRACHYURY on chromatin accessibility at EOMES-bound sites are only weak, and the absence or presence of *Brachyury* does not lead to major changes in *Eomes* regulated gene programs. Thus, in coexpessing cells in the PS during early gastrulation, EOMES suppresses the activation of *Brachyury* regulated PM gene programs to ensure proper lineage specification to AM and DE. This regulation argues against a prevalent concept of the specification of a common mesendoderm progenitor cell type that in successive steps becomes subspecified into different mesoderm and endoderm cell types. Rather pluripotent cells acquire a specific lineage-defining gene program a priori, which is at least in parts regulated by the activities of early Tbx genes. In summary, this study provides a molecular basis for the temporal and spatial order of early generated *Eomes*-dependent lineages which are followed by later-generated and *Brachyury*-dependent lineages that rely on the absence of EOMES.

## Acknowledgements

We thank T. Bass for excellent technical assistance; B.G. Herrmann for providing *T*^2J^ mice, and M. Kyba for the A2lox.Cre ESC line. We are grateful to the Freiburg Galaxy Team (Bioinformatics, University of Freiburg, Germany) and computing support by the state of Baden-Württemberg through bwHPC, University of Tübingen. We acknowledge the Genomics Core Facility at the EMBL (Heidelberg, Germany) for sequencing. The authors acknowledge the support of the Freiburg Galaxy Team: Björn Grüning, Bioinformatics, University of Freiburg (Germany) funded by the Collaborative Research Centre 992 Medical Epigenetics (DFG grant SFB 992/1 2012) and the German Federal Ministry of Education and Research BMBF grant 031 A538A de.NBI-RBC. This study was supported by the German Research Foundation (DFG) through the Heisenberg Program (AR 732/3-1), project grant (AR 732/2-1,) project B07 of CRC 1140 (project ID 246781735), project A03 of CRC 850 (project ID 89986987), project A08 of CRC 992 (project ID 192904750), and Germany’s Excellence Strategy (CIBSS – EXC-2189 – Project ID 390939984) to S.J.A. The work of K.M.S was funded by the EQUIP Program for Medical Scientist, Faculty of Medicine, University Freiburg and by a CRC 992 MEDEP-Fellowship.

## Author contributions

K.M.S., J.W., S.P., A.E.W., L.Z., C.M.S., I.-M.S., and S.J.A. performed experiments. K.M.S., J.W., S.P. and S.J.A. planned and analyzed experiments. K.M.S., J.W., M.T., G.-J.K. and S. performed bioinformatics data analysis. K.M.S, J.W., S.P. and S.J.A. prepared figures, wrote and edited the manuscript with input from all authors, who approved the final manuscript and submission. S.J.A. conceived the study.

## Methods

### Mouse lines and maintenance

*Eomes*^*Δ*Epi^ (*Eomes*^CA/CA^;*Sox2*.Cre) embryos were generated as previously described ^5,35^. *Brachyury* null embryos (*T*^2J/2J^) were a kind gift of B.G. Herrmann. *T*^Cre^ mice, *Eomes*^Cre^ and *Rosa26*^LacZ^ and mice are described previously ^6,30,51^. All mouse strains were maintained on a 129SvEv/C57Bl/6 mixed genetic background. The *Eomes*^*Bra*^ allele was generated using the same strategy as previously described for the GFP knock-in allele ^52^. In brief, *Brachyury*-*V5*.*pA* coding sequence was introduced into the exon 1 start site of the endogenous *Eomes* locus, followed by a removable loxP-flanked neomycin-resistance selection cassette (PGK.Neo).

The 3’ homology arm was flanked by a pMC1.TK negative selection cassette. Targeting vector was electroporated into CCE embryonic stem cells (ESC) and the correctly targeted clones were used to generate germline chimaeras. Embryos were dissected at indicated time points. *Eomes*^*BraΔ*Epi^ embryos were genotyped by PCR using forward primer AGGAAAGGGGCACCTACAATCC and reverse primers GACGCTTTGTCTAAGTCCAGCCTC and GCCTGACACATTTACCTTCAGCAC. Detected WT band has 411 bp, while *Brachyury* knock-in into the *Eomes* locus has 469 bp. For RNA-seq, embryos were genotyped by RT-qPCR for expression of *Eomes* and *Brachyury*. Animals were maintained as approved by the Regierungspräsidium Freiburg (license numbers G11/31 and X19/O2F).

### Cell lines and culture

A2lox.Cre mouse (male) ESCs ^53^ were maintained in Dulbecco’s modified Eagle’s medium (DMEM) containing 15% fetal bovine serum (FBS, Gibco), 2 mM L-glutamine, 1X non-essential amino acids (NEAA), 1 mM sodium-pyruvate, 1X penicillin/streptomycin, 100 µM β-mercaptoethanol, Leukemia inhibitory factor (ESGRO LIF, Merck Millipore, 1000 U/ml), and 2i: CHIR99021 (Axon Medchem, 3 µM) and PD0325901 (Axon Medchem, 1 µM) on a monolayer of mitotically inactivated STO mouse fibroblast cells (SNL76/7) or on 0.1% gelatin-coated dishes at 37°C.

### Generation of Tbx-deficient and of dox-inducible ESCs

Mouse ESCs deficient for *Eomes* and *Brachyury* were generated using TALENs and targeted homologous recombination as previously described ^21^. Resistant colonies were screened by PCR for the integration of the fluorescent reporters and frame-shift deletions. A2lox.Cre mouse ESCs were used to generate dox-inducible gene expression by induced cassette exchange ^53^. Generation of dox-inducible expression of Eomes-GFP (EoGFP) or Brachyury-GFP(BraGFP) was done as previously described ^21^. For dox-inducible expression 5 µg/ml dox was applied for 3 days in differentiation medium. EoGFP and BraGFP constructs contained full-length coding regions of *Eomes* or *Brachyury* fused via a linker to the *Gfp* coding sequence.

### ESC differentiation to mesoderm and definitive endoderm using ActivinA

Prior to differentiation, ESCs were depleted of feeders by splitting for 2-3 passages onto 0.1% gelatin-coated 60 mm dishes. For embryoid body (EB) formation, 200 cells in 40 µl ESGRO Complete Basal medium were grown in ultra-low attachment 96-well plates (Greiner BioOne) for 2 days until EBs had formed. EBs were transferred into 60 mm non-adhesive dishes and differentiated in ESGRO Complete Basal medium with 30 ng/ml human recombinant ActivinA (ActA, R&D systems) for 3 days. For ChIP-seq samples, 2.5×10^5 cells were seeded in 10 ml ESGRO Complete Basal medium into 100 mm non-adhesive dishes. Medium was changed on day 2 and 4 with the addition of 30 ng/ml human recombinant ActivinA and 5µg/ml doxycycline. For CHIR-induced EB differentiation EBs were generated using published protocols^54^ with some modifications, as outlined below. EB formation was performed in ESGRO Complete Basal Medium (Merck Millipore) in the absence of Matrigel. Gastruloids were induced by administration of 3 µM CHIR and, if indicated, with DOX (1 µg/µl, Sigma) for 24 h. In the course of EB culture, the medium was changed at 72h.

### β-Galactosidase staining

Freshly dissected embryos were fixed for 30 min (E7.5) or 60 min (E9.5) at RT using 1 % formaldehyde and 0.2% glutaraldehyde in X-Gal buffer (5 mM EGTA, 0.4 g MgCl_2_ 6H_2_O, 0.4 ml IGEPAL, 0.2 ml deoxycholate in DPBS + Ca/Mg). After washing in PBT 3×5 min, staining was performed overnight at 37°C in 5 mM K-Ferricyanide (K_3_Fe), 5 mM K-Ferrocyanide (K_4_Fe), 0.5 mg/ml X-Gal in X-Gal buffer. Reaction was stopped by washing in PBT and the embryos were post-fixed overnight in 4% PFA in PBS at 4°C. Pictures of whole embryos were taken in 80% Glycerol in PBS. For sectioning embryos were embedded in paraffin and cut into 10 μm thick transversal sections. Sections were counter stained with Eosin to visualize β-Galactosidase non-stained cells. Pictures were taken using Stereo Microscope Leica M165C for whole embryos and Leica DM IL LED for the sections.

### Hematoxylin & Eosin staining

Embryos embedded in paraffin were cut into 10 μm thick transverse sections, deparaffinized and rehydrated using HistoClear, 100%, 96%, 70%, 50% EtOH and H_2_O. Staining was performed in Harris hematoxylin with glacial acetic acid for 3 min and sections were washed in H_2_O. Eosin staining was done for 45 sec. Sections were dehydrated in 50%, 70%, 96% and 100% EtOH and Xylol, covered with a coverslip and mounted using DPX slide mounting medium (Sigma Aldrich, 06522-100ML). Pictures were taken using Microscope Leica M165C for whole embryos and Leica DM IL LED for the sections.

### In situ hybridization

For whole-mount in situ hybridization embryos and EBs were collected in PBT, fixed overnight in 4% PFA in PBS at 4°C and dehydrated in a 25%, 50%, 75% and 3x 100% Methanol in PBT. On the day of the whole mount in situ hybridization embryos were rehydrated in reverse Methanol in PBT series, bleached in 6% H_2_O_2_ in PBT for 30 min (5min for EBs) on ice and washed 3x for 5 min in DEPC-PBT. Embryos were incubated with 10 μg/ml Proteinase K in DEPC-PBT for 5 min (E6.75-7.5) 6 min (E8.5), or 1.6µg/ml Proteinase K in DEPC-PBT for 2min (EBs) and the reaction was stopped using 2 mg/ml glycine in DEPC-PBT for 5 min on ice. Post-fixation was performed on ice in 4 % PFA and 0.2% glutaraldehyde in PBT for 20 min. Next, incubation in prehybridization buffer - 1:1 mixture of DEPC-PBT and hybridization buffer (50% deionized formamide, 5x SSC pH 5.0, 0.05% SDS, 50 μg/ml yeast tRNA, 50 μg/ml Heparin, 60 mM citric acid, 0.1% Tween20 in DEPC-H_2_O) for 10 min at RT in net wells was performed, followed by incubation in the hybridization buffer at 68°C for 2h. The solution was than replaced by pre-warmed hybridization buffer containing 50-100 ng of digoxigenin labelled riboprobe and incubated overnight at 68°C. The next day, 10 min wash in prehybridization buffer, 2x 30 min washes in Wash Buffer I (50 % Formamide, 5x SSC, 1% SDS in DEPC-H_2_O) and 2x 45 min in Wash Buffer II (50 % Formamide, 2x SSC, 0.2% SDS, 0.1% Tween20 in DEPC-H_2_O) were performed at 68°C. Embryos were allowed to cool down to RT and washed 3x 5 min with MAB buffer (150 mM Maleic acid, 100 mM NaCl, 0.1% Tween20, 7.9g/l NaOH, pH 7.5 in H_2_O) for 30 min at RT. For blocking, embryos were incubated in 10% Sheep serum and 2% Blocking Reagent, for nucleic acid hybridization and detection (BBR, Roche, 11096176001) in MAB buffer at RT for 90 min. Antibody solution contained 1 ml of 2% BBR in MAB buffer with embryo powder dissolved at 70°C with agitation, 5 μl of Sheep serum, and 1 μl of Alkaline Phosphatase-linked α-Digoxigenin antibody (Roche, 11093274910) per sample. The solution was nutated at 4 °C for 1h and spun down. The supernatant was diluted to 2.5 ml per sample using 2% BBR in MAB buffer and 20 μl sheep serum per sample. Embryos were incubated in the antibody solution overnight at 4°C. Next day, embryos were extensively washed with MAB buffer. The following day, embryos were washed 3x in freshly prepared NTMT buffer (100 mM Tris-Hydrochloric acid pH9.5, 100 mM NaCl, 50 mM MgCl_2_, 0.1% Tween20 in H_2_O) and color reaction was developed with BM purple (Roche, 1442074) at RT. The reaction was stopped by washing the embryos several times in MAB buffer.

For the double color whole mount in situ hybridization, next to digoxigenin labelled probe, also fluorescein labelled probe was used. First color development was performed using INT-BCIP block solution (Sigma Aldrich, 11681460001) in 0.1 M Tris buffer pH9.5 with 0.05 M MgCl_2_ and 0.1 M NaCl and the pictures were taken. Alkaline Phosphatase was inactivated by washing embryos 3x 5 min in MAB buffer, incubation in MAB buffer at 68°C for 30 min and afterwards with 0.1 M Glycine in PBT pH2.2 for 30 min at RT. To visualize the second riboprobe the protocol was followed from the blocking step until the picture acquisition. Pictures were taken using Stereo Microscope Leica M165FC.

For in situ hybridization of sections, embryos were fixed in the deciduae in 4% PFA/PBS overnight at 4°C and embedded in paraffin. At least three embryos of the same genotype were used. Embryos were cut in 8 μm thick transversal sections, deparaffinized, treated for 5min with 1 μg/ml Proteinase K in PBS, fixed for 15 min and incubated in freshly prepared 0.25% acetic anhydride in TEA-buffer for 10 min. The sections were then incubated in hybridization buffer and following steps were done according to the standard protocol.

### Immunofluorescence staining

Embryos were fixed in 4% PFA overnight at 4°C. The samples were cryo-embedded in 15% Sucrose and 7.5% cold water fish gelatin and cut into 8 μm transversal sections. After permeabilization with 0.2% TritonX-100 in PBS for 20 min at RT and blocking for 2 h with 1% Bovine serum albumin (BSA) in PBS, slides were incubated with primary antibody overnight at 4°C and subsequently with a secondary fluorescence-conjugated antibody and 1:1000 DAPI dilution for 90 min at RT in the dark. The primary antibodies used include anti-TBR2/EOMES (Abcam, ab23345), and anti-V5 (Merk Millipore, ab3792). Secondary antibodies used: anti-Rabbit AlexaFluor 488 or 647 (Thermo Fisher). Experiments were repeated at least three times and comparable images were taken using same excitation intensity, exposure time and gain values. Confocal imaging was performed using the LSM-I-DUO LIVE 510 META Axiovert microscope equipped with a 40x/1.2 C-Apochromat objective W Korr UV-VIS-IR (Carl Zeiss). Excitation of the fluorophores (DAPI, Alexa 488, Alexa 647) was performed with a two-photon laser at 740 nm, and a single photon laser at 488 nm and 633 nm, respectively. Images were processed using FIJI (ImageJ, v2.0) software.

CHIR-induced EBs were fixed in 4% PFA/PBS for 1 h at 4°, permeabilized (0.3% Triton X-100/ PBT, 30 min) and blocked in 1% BSA/PBT for 1 h at room temperature. Primary antibody incubation was performed at 4°C overnight in 1% BSA/PBT, EBs were washed four times in PBT before secondary fluorescence-conjugated antibody incubation was done for 3 h followed by DAPI staining for 30 min at room temperature. Primary antibodies used were anti-brachyury (R&D Systems; AF2085, 1:500), anti-Eomes (abcam, ab23345, 1:500). Secondary anti-goat (A-11056, 1:1000) Alexa Fluor 546-conjugated and anti-rabbit (A-31573, 1:1000) Alexa Fluor 647-conjugated antibodies (both from Thermo Fisher).

### RNA-seq

Total RNA from approximately 25-50 EBs at day 5 of differentiation was isolated using RNeasy Mini kit (Qiagen) and quantified by NanoDrop (Thermo Fisher). Library preparation was either carried out using 0.5 μg of RNA with NEBNext Ultra RNA library Prep Kit for Illumina (New England BioLabs, E7530L) or was performed by Novogene Services, UK, using the Ultra RNA Library Prep Kit. For RNA-seq from E7.5 embryos, the embryos were isolated, and the epiblast and overlying endoderm layer were dissected from extraembryonic portions along the embryonic-abembryonic border. RNA from the epiblasts and overlying endoderm was isolated using the Qiagen RNeasy Micro kit (74004). Libraries were prepared from 10 ng of total RNA from single embryonic samples with the Ovation SoLo RNA-seq Systems kit (NuGEN, 0501-32). All samples were sequenced in four biological replicates from four age-matched individual embryos. Sequencing was performed at Genomics Core Facility (GeneCore, EMBL, Heidelberg, Germany) or at Novogene Services, UK.

### ChIP-seq

ChIP seq was done as described previously ^21^. Briefly, on day 5 of differentiation, EBs were trypsinised and approximately 3×10^7^ cells were used per ChIP. Cells were double cross-linked using 2 mM disuccinimidyl glutarate followed by 1% formaldehyde. Chromatin was isolated, sonicated and incubated with 5µg anti-GFP antibody (Abcam, ab290) coupled to ProteinG beads overnight at 4°C. The next day, beads were washed on ice with RIPA buffer and TBS. DNA was eluted, de-crosslinked and purified. Libraries were prepared with NEBNext Ultra II DNA Library Prep Kit for Illumina (NEB, E7645S), size selected using the SPRI select beads (Beckman Coulter) to obtain fragments between 300-600 bp of size. Samples were sequenced at Novogene Services, UK or at Genomics Core Facility (GeneCore, EMBL, Heidelberg, Germany).

### ATAC-seq

The protocol was modified from the original description of the method ^55^. In brief, EBs at day 5 of differentiation were trypsinized to single-cell suspension, washed with PBS and 50.000 cells were lysed in 50 µl ATAC-seq lysis buffer. Nuclei were mixed with 50 µl of transposition reaction mix containing Tagment DNA Enzyme 1 (Nextera DNA Library Preparation Kit, Illumina). The transposition reaction was carried out at 37°C and 600rpm for 30 min, purified using the Qiagen MinElute Kit and eluted in EB-buffer (Qiagen). DNA was amplified by PCR with an appropriate number of PCR cycles (5-8) using Custom Nextera PCR primer and the NEBNext High-Fidelity 2x PCR Master Mix (Nextera DNA Library Preparation Kit (New England BioLabs, E7530L) and purified with the Qiagen MinElute Kit (Qiagen, 28004). The libraries were size-selected using SPRI select beads (Beckman Coulter) to obtain fragments between <1000 bp of size. DNA concentration was measured with a Qubit Fluorometer (Thermo Fischer, Q32854) and fragment size determined with a Bioanalyzer (Agilent). Libraries were sequenced at Genomics Core Facility (GeneCore, EMBL, Heidelberg, Germany) or at Novogene Services UK.

### ATAC-seq data analysis

Reads were mapped to the genome and differential accessibility analyses were performed using DiffBind tool ^56^ (Galaxy Version 2.10.0) with default settings and FDR threshold of 0.01 and subsequent filtering for minimal fold change (FC) of 2.5 (Galaxy version 1.1.1). Heatmaps were made with ATAC-seq normalized counts to RPKM depicted as the intensity of green color, while the peak profile was created with averaged RPKM values using the computeMatrix and plotHeatmap functions of deepTools (v3.0.2, ^57^). For the visualization in IGV, results from biological duplicates were merged using Merge BAM files (picard v1.56.0), the coverage files were created using bamCoverage (deepTools v3.0.2, ^57^) with bin size 10 bases and normalization to RPKM.

### ChIP-seq data analysis

Reads were mapped to the mm10 genome using Galaxy platform Bowtie2 v2.3.4.3+galaxy0) ^58^ with default settings and the duplicates removed with RmDup ^59^ (Galaxy version 2.0.1). Peaks were detected using MACS2 (v2.1.1.20160309.6, ^60^ with BAMPE format of the input file, default settings for building the shifting model (confidence enrichment ratio against background 5-50; band width 300) and minimum q-value cut-off for detection of 0.01. Peaks were called for each individual replicate and for both replicates together. Intersection (minimal overlap of at least 1bp) of the called peaks using bedtools intersect intervals (Galaxy Version 2.29.2) were defined as high confidence peaks and used for further analysis. EO- and BRA-unique peaks were identified by using the Galaxy DiffBind tool ^56^ (v2.10.0) with default settings and a minimal FDR threshold of 0.01 and subsequent filtering for minimal fold change (FC) of 2.5 (Galaxy version 1.1.1). EOMES and BRACHYURY common peaks were defined by minimal overlap of 1bp of EoGFP and BraGFP high confidence peaks using bedtools intersect intervals (Galaxy Version 2.29.2) followed by subtraction of EO- and BRA-unique peaks (Galaxy version 2.27.1). Differential binding of EoGFP and BraGFP in sKO versus dKO was analyzed by using the Galaxy DiffBind tool ^56^ (v2.10.0) with default settings and a minimal FDR threshold of 0.01 and subsequent filtering for minimal fold change (FC) of 2.5 (Galaxy version 1.1.1). For visualization in IGV, results from biological duplicates were merged using Merge BAM files (Galaxy version 1.2.0), the coverage files were created using bamCoverage (Galaxy version 3.3.2.0.0, ^57^) with paired-end extension, bin size 10 bases and normalization to RPKM. Heatmaps were made with ChIP-seq normalized counts to RPKM depicted as the intensity of blue color, while the peak profile was created with averaged RPKM values using the computeMatrix and plotHeatmap functions of deepTools (Galaxy version 3.3.2.0.0 and 3.3.2.0.1, respectively, ^57^).

### RNA-seq data analysis

#### RNAseq of embryos

Sequenced reads were mapped to and annotated by using the mouse reference genome GRCm38/mm10 (iGenomes, Illumina; archive-2015-07-17-32-40 and archive-2015-07-17-33-26 for GRCM38 and mm10, respectively) containing the chromosomes 1-19, X, Y and M using Rsubread v1.28.1 package in R v3.4.3 ^61^. Genomic features were counted using the Rsubread::featureCounts function ^61^. Differential expression analysis was carried out using DESeq2 v1.18.1 in R ^62^, which applies Negative Binomial GLM fitting and Wald statistics on the count data, with subsequent filtering for adjusted p-value<0.05 and log_2_FC as indicated in the figure legends. Prior to fitting and statistical analysis, counts were normalized by library size (DESeq2::estimateSizeFactors) and gene-wise dispersion (DESeq2::estimateDispersions) to normalize the gene counts by gene-wise geometric mean over samples. For plotting the heatmaps, gene-wise scaling was performed on the normalized counts to cluster the counts for the expression tendency between the samples (base::scale function). Scaling centers the count values, by setting the gene-wise mean to zero, which explains why some count values are negative (below gene-wise mean). Visualization of the clustered data was performed using the pheatmap::pheatmap function ^63^ (pheatmap v1.0.10) with deactivated function-intrinsic clustering.

For the visualization of Volcano plots, Venn diagrams and coverage tracks in the Integrated Genome Viewer (IGV) v2.3.93 ^64^, mapping was performed on Galaxy platform ^65^ using Bowtie2 (v2.3.4, ^58^) on mm10 reference genome with default settings. For IGV coverage tracks obtained bam files from biological triplicates were merged using Merge BAM files tool in Galaxy (picard v1.56.0) and the duplicates were removed by RmDup tool ^59^ (samtools v1.3.1). Coverage files were created using bamCoverage tool (deepTools v3.0.2, ^57^) with bin size 10 bases and reads per kilobase per million (RPKM) normalization. Volcano plots were generated using Volcano plot tool in Galaxy (v0.0.3) with a cut off p-value<0.05 and log_2_FC as indicated in the figure legends.

#### RNAseq of EBs

Low-quality bases and adapter-containing reads were trimmed using Galaxy platform Trim Galore ^66^ (Galaxy Version 0.4.3.1) and reads were aligned to the mouse reference genome mm10 (mm10_UCSC_07_15, RNA STAR ^67^ Galaxy Version 2.7.2b) with default parameters. Duplicates were removed with RmDup ^59^ (Galaxy version 2.0.1) and genomic features were counted using htseq-count (Galaxy Version 0.9.1) with following settings: mode-union, stranded-no, minimum alignemt quality-10 and feature type-exon. Differential expression analysis was carried out using DESeq2 (Galaxy Version 2.11.40.6+galaxy1) with subsequent filtering for adjusted p-value<0.05 and log_2_FC as indicated in the figure legends

#### Genomic Regions Enrichment of Annotations

Statistical enrichment for association between genomic regions defined as EOMES and BRACHYURY common or unique peaks and annotations of putative target genes was performed using Genome Regions Enrichment Annotations Tool (GREAT) online tool (v4.04 ^37^) using basal plus extension setting. Each annotated gene was assigned a basal regulatory region of 5 kb upstream and 1 kb downstream of the transcription start site, and the regulatory domain was extended in both directions up to 1,000 kb until the nearest gene’s basal domain.

#### Genomic distribution pies

Peak annotation and visualization of genomic locations of EO-unique, BRA-unique and EO+BRA common peaks were performed using ChIPseeker tool ^68^ in Galaxy v1.18.0+galaxy1 with default settings.

#### Gene Ontology overrepresentation

For GO-term analysis, The Database for Annotation, Visualization and Integrated Discovery (DAVID) v6.8 ^69^ was used. Representative GO-terms are shown using threshold count 5, EASE 0.1, and Benjamini p-value adjustment, by sorting for adjusted p-value<0.05 and highest fold change.

#### Motif enrichment analysis

For motif discovery and motif enrichment analysis, we used respectively: Meme Suite ^70^ (v5.1.1) with the settings: classic motif discovery mode, zero or one occurrence per sequence (zoops), top 5 motifs, min width 6, max width 50 and Homer ^40^ (v4.9.1) 80 findMotifsGenome.pl on peaks summits +/-75bp based on MACS2(v1.4.2) narrowPeak and summits in bed files. For motif quantification against background and peak loci, the R packages Biostrings ^71^ (v2.62.0) and universalmotif ^72^ (v1.12.1) were used to match the position weight matrices along with their prior background base probabilities from the above Homer and Meme motifs via the matchPWM function, using a minimum scoring threshold of 85% to fit motifs on all forward and reverse reading frames on both the peak loci and the UCSC Mus Musculus mm10 genome provided by the BSgenome package ^73^ (v1.62.0).

#### Clustering and visualization of mouse gastrulation atlas data

Processed atlas data on mouse organogenesis from ^28^ were downloaded from ArrayExpress (accession number: E-MTAB-6967). The following time points and sequencing batches were analyzed: E6.5 (sequencing batch 1), E6.75 (sequencing batch 1), E7 (sequencing batches 1, 2 and 3), E7.25 (sequencing batch 2) and E7.5 (sequencing batches 1 and 2), E8 (sequencing batches 2 and 3), E8.25 (sequencing batch 2) and E8.5 (sequencing batches 2 and 3). Cells defined as doublets in the study were removed from the analysis. Integration of datasets from different time points and sequencing batches was performed using Harmony package in Seurat version 3 with default settings ^74^. Ribosomal genes (small and large subunits), as well as genes with Gm-identifiers were excluded from the data before integration. The integrated dataset contained 115,713 cells. A focused analysis of cells expressing *Eomes* or *Brachyury* which includes *Eomes* single positive, cells, *Brachyury* single positive cells and *Eomes/Brachyury* double positive cells at E7 (sequencing batch 3), E7.5 (sequencing batch 2) and E8 (sequencing batch 3) was performed using VarID ^29^. Identical settings were used for VarID analysis at each timepoint. Cells with a total number of transcripts of less than 6000 were discarded, and count data of the remaining cells were normalized by downscaling. Cells having normalized *Eomes* or *Brachyury* transcript counts of more than 0.3 were further analyzed (6,914 cells at E7, 3,746 cells at E7.5, and 3,766 cells at E8) and further clustered and visualized using VarID with the following parameters: large=TRUE, pcaComp=100, regNB=TRUE, batch=batch, knn=50, alpha=10 and no_cores=20. Dimensionality reduction of the datasets was performed using UMAP.

#### Quantification and statistical analysis

All RNA-seq experiments were performed as 3 biologically independent experiments, ATAC-seq and ChIP-seq as 2 biologically independent experiments. For *in situ* hybridization experiments with mouse embryos at least 3 embryos of different genotype were used for the same developmental stage, and the experiment was repeated 2 times with similar results. Immunofluorescence stainings were performed at least 2-3 times and the representative images are shown. For *in situ* hybridization experiments and immunofluorescence stainings with gastruloids at least 6 gastruloids of different condition were used for the same timepoint with similar results and the representative images are shown.

#### Data availability

RNA-seq, ATAC-seq, and ChIP-seq data of embryos and cells deficient for *Eomes* and *Brachyury* (Tosic et al. 2019) that support the findings of this study have been deposited in the Gene Expression Omnibus (GEO) under accession code GSE128466. Newly generated data will be available in the GEO database upon publication.

## Figure titles and legends

**Figure S1.**
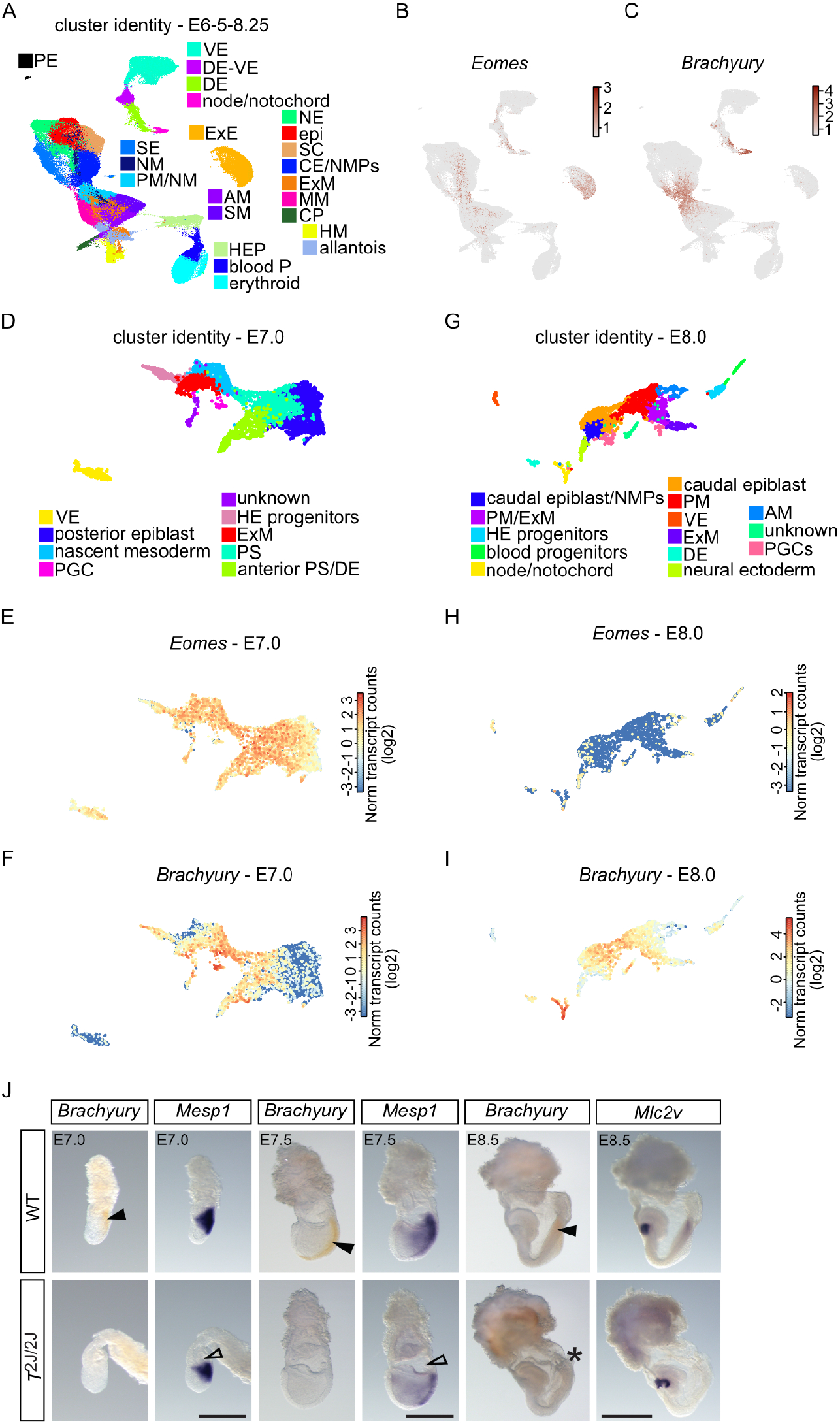
*Eomes* and *Brachyury* specify distinct cell types at different gastrulation stages. **A-C)** UMAPs of total cell composite from E6.5 – E8.25 embryos with assigned identities to different clusters (A), and UMAPs showing expression of *Eomes* (B) or *Brachyury* (C) (data from ^28^). Scale bars represent log2 normalized transcript counts. PE – parietal endoderm, VE visceral endoderm, DE-VE – definitive and visceral endoderm, DE – definitive endoderm, SE – surface ectoderm, NM – nascent mesoderm, PM/NM – posterior and nascent mesoderm, AM – anterior mesoderm, SM – somitic mesoderm, ExE – extraembryonic ectoderm, NE – neuroectoderm, Epi – epiblast, SC – spinal cord, CE/NMPs – caudal epiblast/neuromesodermal progenitors, ExM – extraembryonic mesoderm, MM – mixed mesoderm, CP – cardiac progenitors, HM – head mesenchyme, HEP – haematoendothelial progenitors, P – progenitors. **D - I)** UMAPs of extracted *Eomes* and/or *Brachyury*-expressing cells from (D-F) E7.0 and (G-I) E8.0 embryos showing (D, G) assigned cell identities to different clusters, and expression levels of (E, H) *Eomes* or (F, I) *Brachyury* (data from ^28^). Scale bars represent log2 normalized transcript counts. Extraembryonic ectoderm was excluded from UMAPs. VE – visceral endoderm, PGC – primordial germ cell, HE – haematoendothelium, ExM – extraembryonic mesoderm, PS – primitive streak, DE – definitive endoderm, NMP – neuromesodermal progenitors, PM – posterior mesoderm, AM – anterior mesoderm. **J)** Double whole mount *in situ* hybridization of WT and *T*^2J/2J^embryos at indicated embryonic timepoints for *Brachyury* (in orange, filled arrowheads) and cardiac mesoderm markers *Mesp1* and *Mlc2v* (in blue, open arrowheads) showing that heart mesoderm is normally specified in *T*^2J/2J^ embryos. Scale bars 500 μm. Asterisk indicates paucity of most posterior mesoderm/allantois in E8.5 *T*^2J/2J^ embryos.

**Figure S2.**
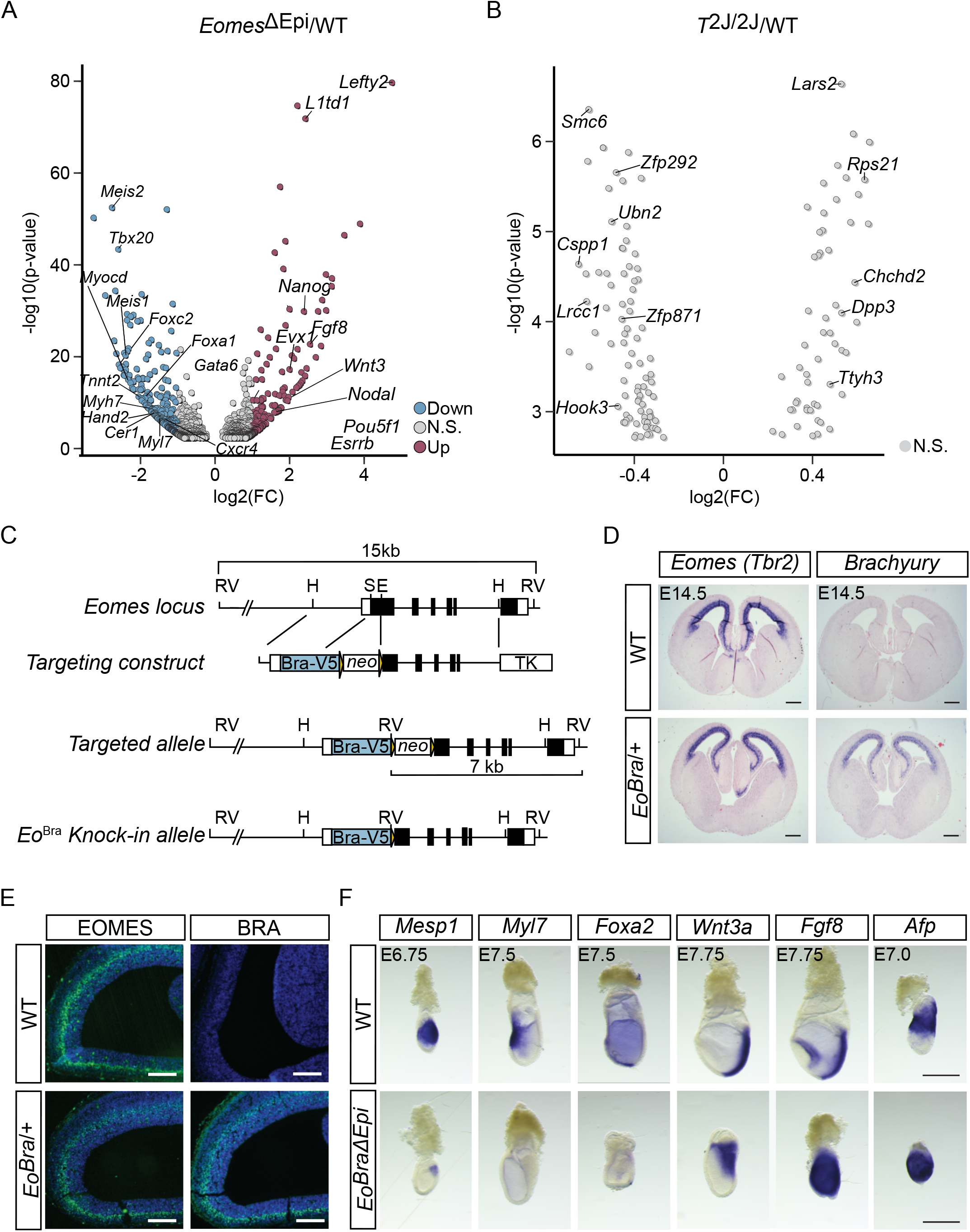
*Brachyury* does not functionally compensate for the lack of *Eomes* in the epiblast. **A)** Volcano plot of differentially expressed genes in *Eomes*^ΔEpi^ compared to WT embryos, adjusted p-value<0.05; log2(FC)>1.0 for upregulated genes, log2(FC)<-1.0 for downregulated genes. **B)** Volcano plot showing no significantly differentially expressed genes in *T*^2J/2J^ compared to WT embryos at E7.5, adjusted p-value<0.05; log2(FC)>1.0 for upregulated genes, log2(FC)<-1.0 for downregulated genes. **C)** Schematic of the targeting strategy to insert *Brachyury*-*V5* (*Bra-V5*) coding sequence into the first exon of *Eomes* coding region to replace *Eomes* with *Brachyury* expression. RV - EcoRV; H - HindIII; S – SphI; E – EagI, neo – neomycin resistance cassette; TK – thymidine kinase; yellow triangles – LoxP sites. **D)** *In situ* hybridization of E14.5 mouse brain sections showing *Brachyury* mRNA expression in the cerebral cortex of *Eomes*^Bra/+^ heterozygous embryos, resembling the expression of *Eomes/Tbr2* in intermediate cortical progenitor cells. *Brachyury* expression is normally absent in the cerebral cortex of WT embryos. Scale bars 500 μm. **E)** Immunofluorescence staining of BRACHURY-V5 protein in the cerebral cortex of E14.5 *Eomes*^Bra/+^ embryos, which is absent in the WT. For comparison IF for endogenous EOMES (TBR2) indicates expression in intermediate progenitor cells of the subventricular zone in both WT and *Eomes*^Bra/+^ cortices. Scale bars 500 μm. **F)** Whole mount *in situ* hybridization of E6.75 - E7.75 WT and *Eomes*^BraΔEpi^ embryos shows absence of cardiac markers *Mesp1* and *Myl7*, absence of the DE marker *Foxa2*, upregulation of putative BRACHYURY targets *Wnt3a* and *Fgf8*, and persistence of the VE marker *Afp* in the *Eomes*^BraΔEpi^ embryos compared to WT controls. Scale bars 200 μm.

**Figure S3.**
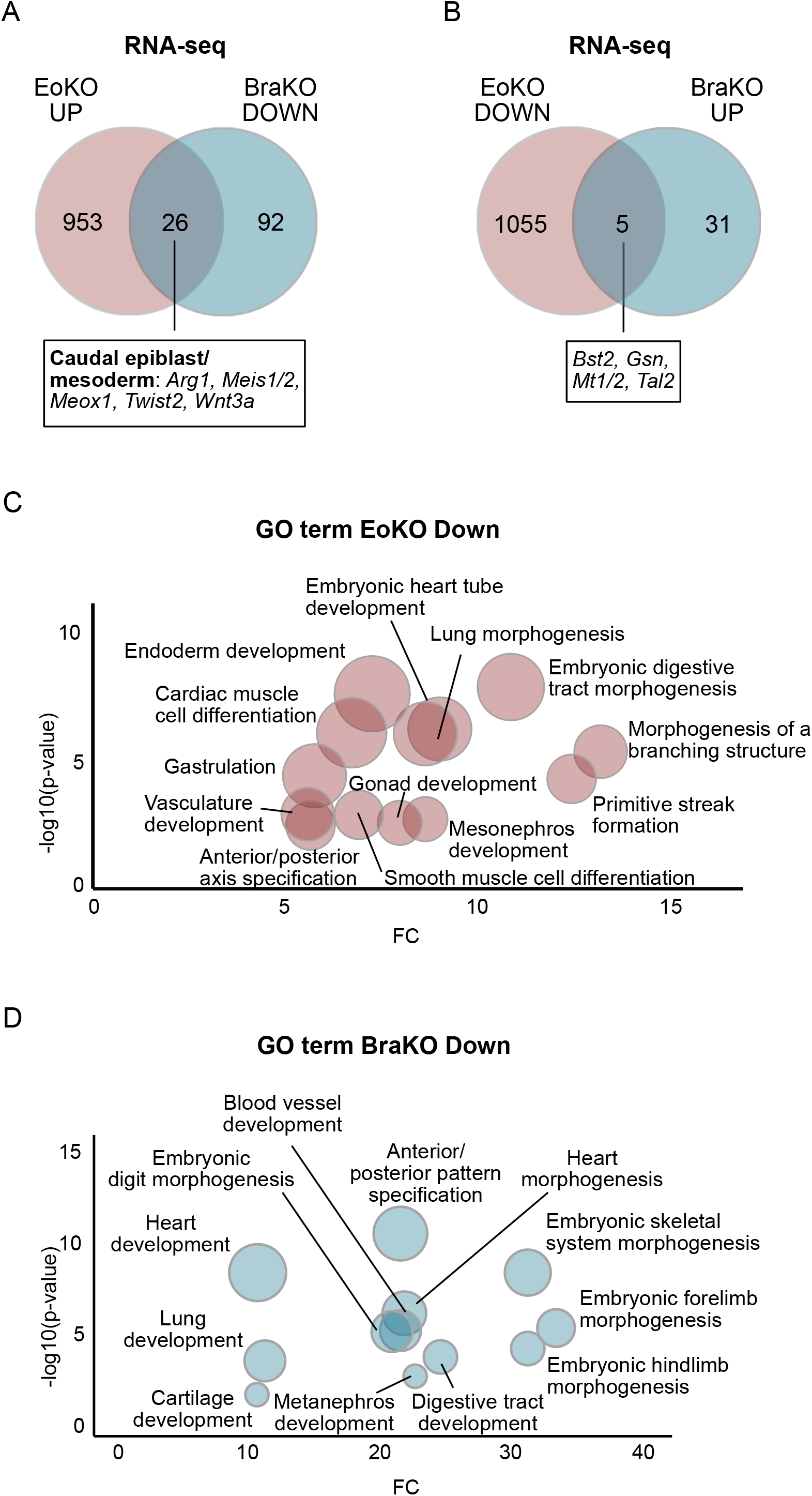
Sets of specific *Eomes* and *Brachyury* target genes and programs. **A, B)** Venn diagram of RNA-seq analyses showing the overlap of genes upregulated in EoKO and downregulated in BraKO compared to WT EBs (A) and vice versa (B). *Brachyury*-target genes are upregulated in EoKO EBs. **C, D)** Gene ontology (GO) term enrichment analysis of genes downregulated in EoKO (C) or BraKO (D) compared to WT. Circle size is proportional to the number of genes (minimum 5).

**Figure S4.**
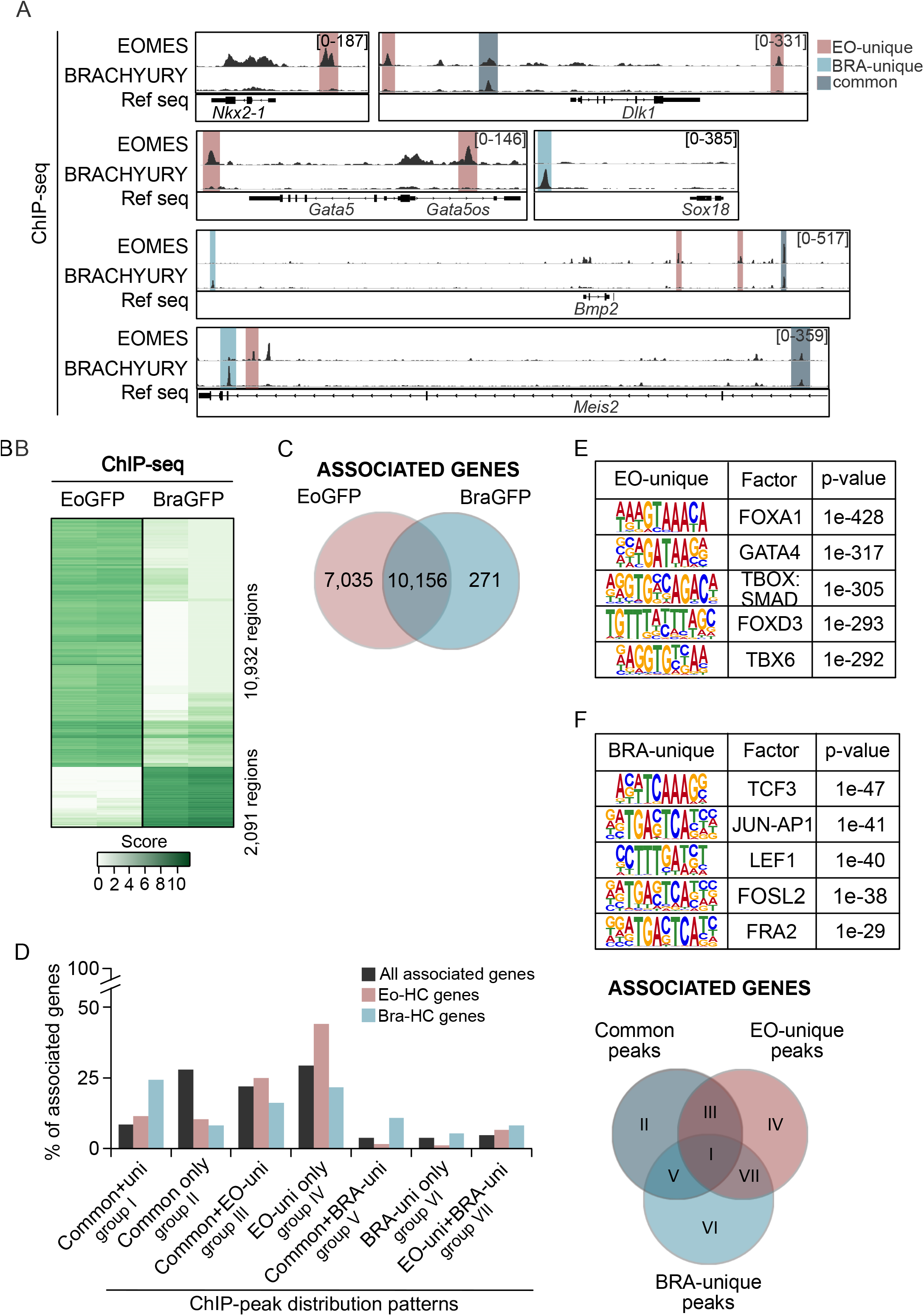
EOMES and BRACHYURY show complex and largely non-overlapping binding patterns at target genes. **A)** Examples of ChIP-seq coverage tracks of EOMES- and/or BRACHYURY-bound regions close to target genes (*Nkx2-1, Dlk1, Gata5(os), Sox18, Bmp2, Meis2*) in ActivinA-induced EBs (EoKO+EoGFP and BraKO+BraGFP). Counts normalized to RPKM are indicated. **B)** Binding affinity heatmaps of differentially bound regions comparing EOMES-GFP (EoGFP) with BRACHYURY-GFP (BraGFP) in differentiating EBs (EoKO+EoGFP and BraKO+BraGFP) analyzed by ChIP-seq of two independent experiments. **C)** Venn diagram showing the large overlap of associated genes (numbers indicated) to EOMES-GFP and BRACHYURY-GFP ChIP-seq binding peaks in differentiating EBs. **D)** Bar graphs show the frequency of genes associated to the different patterns of EOMES- and BRA-binding within all target genes, *Eomes* (Eo-HC), or *Brachyury* (Bra-HC) high-confidence target genes. The percentage is depicted as ratio of overlap of associated genes with all, Eo-HC and Bra-HC genes to the total number of all, Eo-HC and Bra-HC genes. Venn diagram is identical to Figure 3C using different labels to identify groups of genes associated to ChIP-peak distributions in the bar graph diagram. **E, F)** Representative transcription factor-binding motif enrichment within EO-unique over BRA-unique binding sites and *vice versa*. Prevalent Tbx-binding motifs were not included. P-values are indicated.

**Figure S5.**
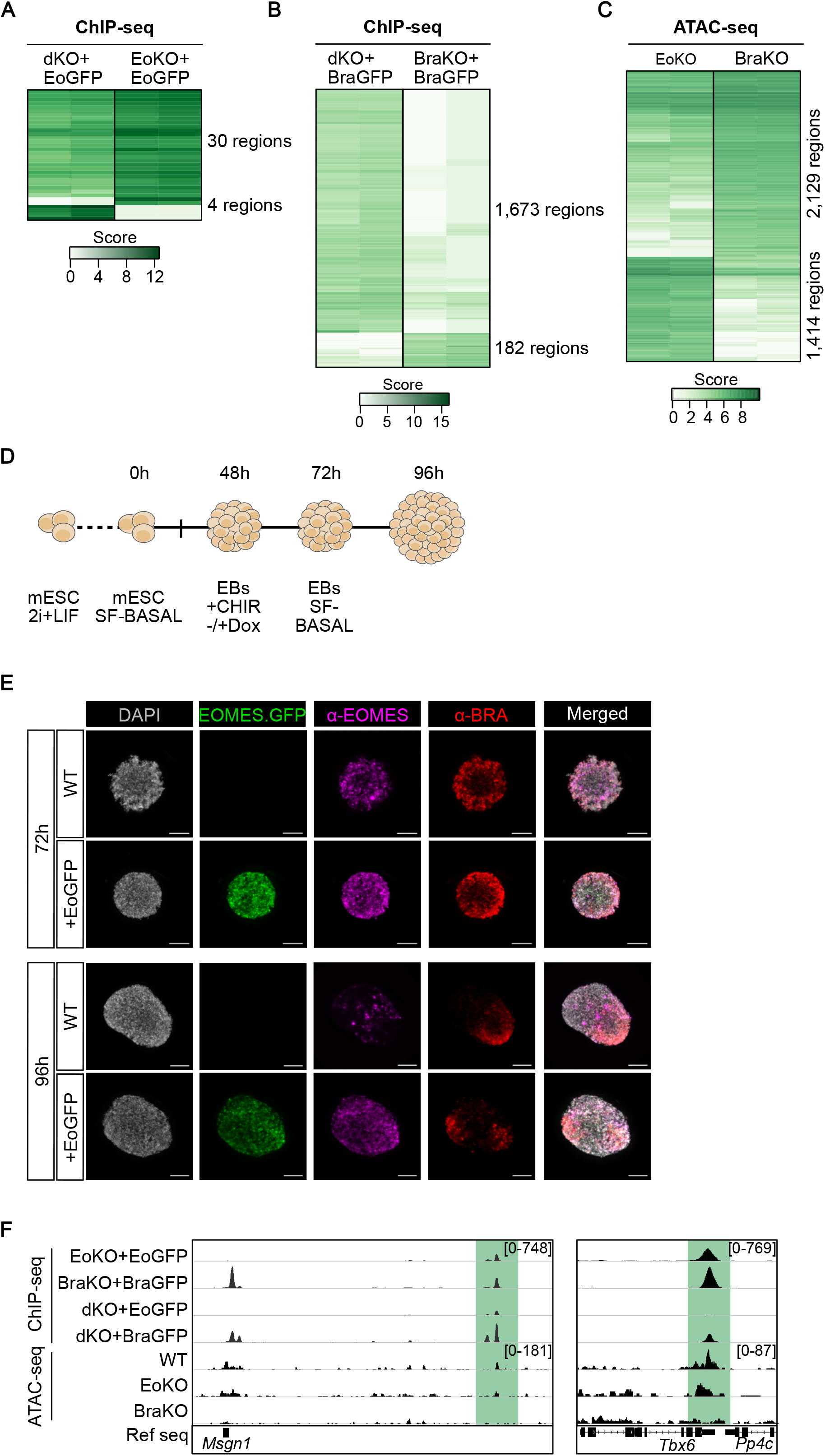
EOMES restricts BRACHYURY functions and competing gene programs. **A)** Binding affinity heatmap of differentially bound regions of EOMES-GFP in dKO versus EoKO differentiating EBs identified by ChIP-seq shows that EOMES binding is almost unaffected by the presence or absence of BRACHYURY, since only 34 differentially bound regions were found. **B)** Binding affinity heatmap of differentially bound regions of BRACHYURY-GFP in dKO versus BraKO differentiating EBs shows that BRACHYURY binding is largely enhanced in the absence of EOMES in dKO cells, indicated by 1,673 regions with increased binding. **C)** Binding affinity heatmap of differentially accessible regions (DARs) between EoKO and BraKO differentiating EBs analyzed by ATAC-seq of two independent experiments. **D)** Schematic illustration of CHIR-induced embryoid body (EB) differentiation. mESC – mouse embryonic stem cells; SF – serum free; h – hour of differentiation; Dox – doxycycline. **E)** Immunofluorescence staining of EOMES and BRACHYURY protein in CHIR-induced EBs at indicated timepoints. *Eomes-GFP* expression is induced in WT+EoGFP EBs with no change in *Brachyury* expression. Scale bars 100 μm. **F)** Examples of ChIP-seq coverage tracks of EOMES- and/or BRACHYURY-bound region in ActvinA-induced EBs (EoKO+EoGFP and BraKO+BraGFP or dKO+EoGFP and dKO+BraGFP) close to *Brachyury*-dependent target genes (*Msgn1* and *Tbx6*). *Brachyury*-target genes show preferential BRACHYURY binding in absence of EOMES and loss of open chromatin in BraKO compared to EoKO and WT as shown by ATAC-seq coverage tracks of WT, EoKO and BraKO ActivinA-induced-EBs. Counts normalized to RPKM are indicated.

